# Drugging Disordered Proteins by Conformational Selection to Inform Therapeutic Intervention

**DOI:** 10.1101/2024.07.03.601611

**Authors:** Bryan A. Bogin, Zachary A. Levine

## Abstract

Drugging intrinsically disordered proteins (IDPs) has historically been a major challenge due to their lack of stable binding sites, conformational heterogeneity, and rapid ability to self-associate or bind non-specific neighbors. Furthermore, it is unclear whether binders of disordered proteins i) induce entirely new conformations or ii) target transient pre-structured conformations via stabilizing existing states. To distinguish between these two mechanisms, we utilize molecular dynamics simulations to induce structured conformations in islet amyloid polypeptide (IAPP), a disordered endocrine peptide implicated in Type II Diabetes. Using umbrella sampling, we measure *conformation-specific* affinities of molecules previously shown to bind IAPP to determine if they can discriminate between two distinct IAPP conformations (fixed in either ⍺-helix or β-sheet). We show our two-state model of IAPP faithfully predicts the experimentally observed selectivity of two classes of IAPP binders while revealing differences in their molecular mechanisms of binding. Specifically, the binding preferences of foldamers designed for human IAPP was not fully accounted for by conformational selection, unlike β-breaking peptides designed to mimic IAPP self-assembly sequences. Furthermore, the binding of these foldamers, but not β-breaking peptides, was disrupted by changes in the rat IAPP sequence. Taken together, our data quantifies the sequence and conformational specificity for IAPP binders and reveals conformational selection sometimes overrides sequence-level specificity. This work highlights the important role of conformational selection in stabilizing IDPs, and it reveals how fixed conformations can provide a tractable target for developing disordered protein binders.

## INTRODUCTION

Intrinsically disordered proteins (IDPs) are widespread in all living organisms and make-up around one-third of the human proteome.^1^ In contrast to folded proteins, IDPs do not fold into sets of well-defined structures, but rapidly sample an ensemble of conformations.^2^ This conformational diversity enables IDPs to carry out multiple roles as transcription factors, chaperones, and hubs of signaling networks with a high sensitivity to minor changes in pH, temperature, or concentration.^2,3^ The regulation of IDP conformations in biology is tightly controlled, and dysfunction of IDPs are implicated in some of the most abundant human diseases, including Alzheimer’s, Parkinson’s, Diabetes, and cancers.^4^ Unfortunately, the same structural diversity that enables IDPs to be multifunctional creates additional challenges towards their druggability, such as how to target pathological conformations while leaving functional conformations alone.^2^ Despite continual improvements of traditional biophysical techniques such as Nuclear Magnetic Resonance (NMR) or single-molecule Förster Resonance Energy Transfer (smFRET) to capture dynamic information encoded in IDPs, the ability to discern between functional and pathological states within large ensembles remains extremely challenging.^2,5^ Thus, new strategies are required to develop drugs for IDPs, distinguish between distinct sub-populations of their conformations, and modulate their functions within the cell.

Despite these challenges, several groups have successfully used natural or synthetic compounds to bias the conformations of IDPs by changing their chemical environment.^6,7^ Previous work which demonstrated that natural cosolvents and osmolytes can rearrange the patterning of IDP stickers and spacers has proven to be a powerful tool to globally denature or stabilize disordered peptides.^6^ However, this binding is largely non-specific, and denaturing compounds are generally unsuitable for therapeutic applications.^8,6^ Alternatively, synthetic peptide-mimetics that recapitulate the folding of proteins (termed as ‘foldamers’) have considerable potential to generate chemically diverse and tunable IDP binders.^2,9,10^ Recently, α-helix mimicking foldamers have been successful at preventing the pathological aggregation of islet amyloid polypeptide (IAPP)^11^, α-synuclein^12^, amyloid beta (Aβ)^13^, and p53^14^. In the case of IAPP, foldamers containing a synthetic oligoquinoline backbone were threaded with diverse chemical moieties and incubated with IAPP in order to stabilize alpha-helical intermediates, preventing the pathological conformational switching and aggregation of the peptide.^15^ Notably, these IAPP-targeted foldamers blocked the formation of toxic macromolecular assemblies and preserved the integrity of cellular membranes exposed to pathological amounts of IAPP.^16^ Lastly, many groups employ rational design strategies based on experimental structures of IDP aggregates to bind and inhibit the aggregation of larger protein assemblies.^17,18,19,20^ One example is a beta-breaking modifications to a 6-residue peptide derived from the aggregation-prone ANFLVH region of IAPP significantly reduces the toxicity of pathological IAPP oligomers.^17^ This example is especially notable since only a small methyl group (α-aminoisobutyric acid - AIB) was used to convert a highly aggregating peptide into an aggregation-inhibitor capable of preventing IAPP toxicity by reducing pathological β-sheet formations.^17,21^

While molecular foldamers and β-inhibitors show great promise as IDP regulators through their preference for a single secondary structure, alpha helices and β-sheets, respectively), their mechanism of action is still unclear.^17,22^ To elucidate their mechanisms, we turn to all-atom molecular dynamics (aa-MD) simulations, which can predict binding affinities of molecules across multiple conformations that are currently inaccessible by experimental methods.^23^ We chose to use IAPP as our target IDP since both foldamers and β-inhibitors have been developed for it.^16,15,20,24,25^ Additionally, some of these IAPP binders have been biophysically characterized, allowing us to benchmark our models predictions to experimental results.^17,22^ IAPP undergoes toxic conformational switching that precedes aggregation, membrane-leakage, and β-cell damage thought to contribute to Type II Diabetes (T2D).^16,24,26,27^ However, the exact connection between its aggregation propensity and biological toxicity remains unclear. Small-molecules able to bind only certain IAPP conformations would not only be helpful in connecting its toxic conformational switching to T2D, but also towards developing therapeutics to prevent its gain-of-toxic function.

In this work, we propose a novel computational strategy for predicting conformation-selective inhibitors of IDPs using aa-MD simulations. This strategy relies on rationally designing binders for specific transient conformations of interest (Figure 1A). Using IAPP as a proof of concept, we asked whether high-affinity IAPP binders prefer specific conformations or short linear sequence motifs (Figure 1B). We show that a simple two-state fixed-conformational model of IAPP is sufficient to recapitulate the conformational binding preferences of high-affinity ligands. Further, we quantify the relative conformational and sequence selection of foldamers and β-inhibitors, which are not accessible by experimental measurements—but inform their different mechanisms of action. Lastly, we compare the human IAPP sequence with its rat IAPP ortholog to determine the extent to which they disrupt the binding affinities of IAPP binders to specific linear sequences. By making comparisons between IAPP inhibitors, non-inhibitors, and non-binder controls, we reveal the degree to which IAPP binders exhibit conformational selection and the extent to which they predict the stabilization of that conformation (Figure 1C).

**Figure 1.**
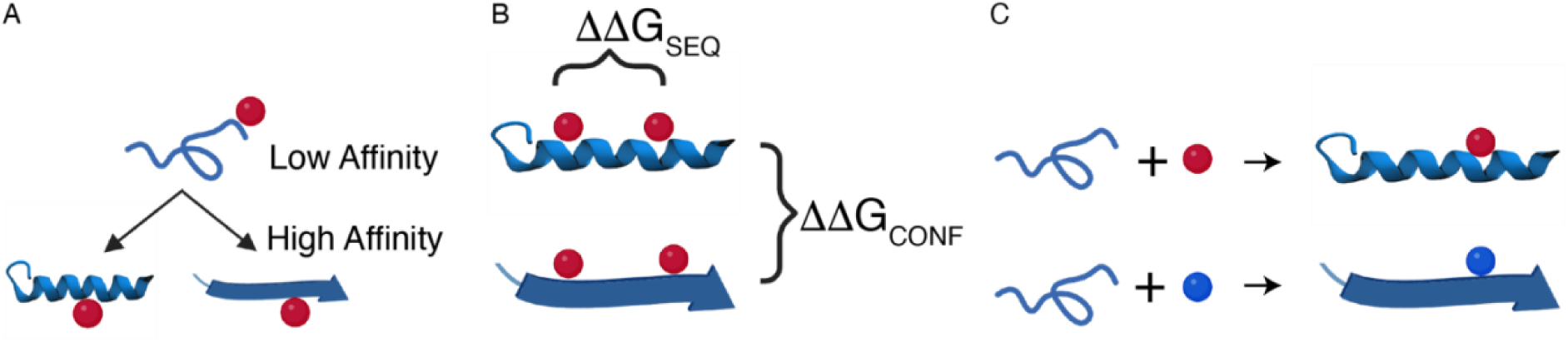
**(A)** Stabilizing IDP conformations of interest by designing high-affinity ligands for target conformations. **(B)** In silico prediction of binding affinities to measure conformational (ΔΔG_CONF_) and sequence selection (ΔΔG_SEQ_) across an IDP sequence. **(C)** Conformation-specific molecules can stabilize transient IDP conformations and avoid pathological ones.

## SIMULATION METHODS

### Theoretical Modeling

Atomistic simulations were conducted using the GROMACS 2022.4 package.^28^ CHARMM36^29^ (updated July 2022) and TIP3^30^ were used for the protein and water forcefields, respectively. CGenFF^31^ (https://cgenff.umaryland.edu/) was used to generate forcefield parameters for the small molecule foldamers.^22^ For the β-Inhibitor (seq XNFXVH) and control peptide (seq ANFLVH), CHARMM36^29^ was used for the forcefield, which already includes the parameters for 2-Aminoisobutyric acid (AIB or X).^17^ Helical and beta strand all-atom peptides of N-terminal human IAPP (seq KCNTATCATQRLANFLVHSSNN), C-terminal human IAPP (seq SSNNFGAILSSTNVGSNTY), rat N-terminal IAPP (seq KCNTATCATQRLANFLVRSSNN), and rat C-terminal IAPP (seq SSNNLGPVLPPTNVGSNTY) were built in Avogadro using the peptide builder tool.^32^ The disulfide bond was added between residues 2 and 7 for the human and rat N-terminal peptides, as observed experimentally.^33^ The histidine residue in human IAPP N-terminal peptides were neutralized (by the addition of a proton to the δ-nitrogen, ND1). A leapfrog algorithm was used to integrate Newton’s equations of motion with a 2 fs timestep. Short-range van der Waals and Coulomb potential cut-offs were set to 1.2 nm. Long-range electrostatics were calculated using the Particle Mesh Ewald (PME) algorithm^34^, which increases computational efficiency of the calculations using Fourier transforms. Periodic-boundary conditions were used in all directions to minimize boundary effects.

### Protein and Small Molecule Preparation

In total, 48 systems were prepared in a 10 nm x 10 nm x 10 nm cubic box containing a combination of IAPP targets (human IAPP targets:_N-terminus_beta-strand, C-terminus_beta-strand, C-terminus_helical, N-terminus_helical; rat IAPP targets: N-terminus_beta-strand, C-terminus_beta-strand, N-terminus_helical, C-terminus_helical) paired with a distinct IDP binder (β-Inhibitor^17^, control peptide^17^, right-handed IAPP foldamer^15^, left-handed IAPP foldamer^22^, right-handed amyloid-beta (Aβ) foldamer^35^, and a control poly-A peptide). N-terminal IAPP peptides were given C-terminal amidation caps to avoid terminal charge interactions. C-terminal IAPP peptides were given N-terminal acetylation and C-terminal amidation caps to recapitulate the physiological capping of IAPP, which is only C-terminally amidated.^36^ The β-Inhibitor^17^, control peptide^17^, and PolyA control were C-terminally amidated. The foldamers^15,22^ were unmodified from Miranker et al.^22^, but were folded into either right-handed or left-handed conformations in prior simulations (Figure S1A). The IAPP backbone was first aligned along the x-axis and then constrained in position by harmonic restraints using a force of 1000 KJ/mol-nm^2^. IDP binders were then randomly positioned within the periodic box, which differed in each technical replicate. Since most of complexes were charged, sodium and chloride counter ions were added to neutralize the net charge of the system and provide a physiological ionic strength of 150 mM. Each configuration was then energy minimized to remove steric clashes before the start of molecular dynamics using two distinct steepest decent simulations—the first simulation with no bond constraints followed by a second simulation where only hydrogen bonds were constrained.

### Deriving Optimal IAPP-Binder Complexes for PMF Calculations

Energy minimized IAPP conformations (fully α-helical or beta-stranded) plus one of the six binders were used as the starting configuration for subsequent equilibration simulations designed to capture stable heterodimers. For each of the 48 IAPP-binder pairs, four technical replicates were constructed with identical starting conditions except for the initial randomized placement of the binder within the periodic box, which resulted in a total of 192 equilibration simulations (Figure S1C). Each equilibration simulation was run for 200 ns under an NVT ensemble (constant number of atoms, volume, and temperature) at a temperature of 300 K using the modified velocity rescaled (v-rescale) thermostat, which accurately recapitulates a canonical thermodynamic ensemble.^37^ This resulted in over 38.4 μs of equilibration simulations alone. During binding, harmonic restraints of 1000 KJ/mol-nm^2^ were used to keep IAPP Cα atoms restrained to ensure that protein conformations did not change during equilibration. Complexation of the binder to IAPP was monitored using the center-of-mass distance between the binder and IAPP backbone. Since binding and unbinding occurred frequently throughout the 200 ns (Figure S2B), the final binding site was obtained using the gromos clustering algorithm (0.2 nm cutoff)^28^ by selecting the most populous complex (Figure S1C). We then compared the four replicates and selected the most likely binding site to IAPP based on if most replicates recapitulated binding within ±2 amino acids of one another. For complexes that did not meet this criterion (i.e. equal binding to different IAPP locations), the complex with the largest amount of time occupied by the binder was utilized as the starting configuration for subsequent PMF calculations (*see Attachment 1*).

### Potential of Mean Force Simulations

To determine the binding energy of each complex we ran umbrella sampling on each of the final configurations derived from equilibration simulations. A harmonic potential with a spring constant of 5000 KJ/mol-nm^2^ was used to pull the binder roughly 3 nm away from its bound position on IAPP at a constant velocity of 0.0002 nm/ps. For completeness, binders were also sometimes pushed 1 nm towards IAPP to ensure a large repulsive force and to confirm that starting configurations took place at the minimum of the Lennard-Jones potential (Figure S4A). Since the IAPP backbone was always aligned along the x-axis, the pulling reaction coordinate was always orthogonal (+y, −y, +z, or −z) depending on the cartesian coordinate closest to the normal vector along the binders center-of-mass. Two additional harmonic potentials (with spring constants of 5000 KJ/mol-nm^2^) were applied to each of the remaining dimensions to restrict diffusion outside of the pulled dimension. Along each one-dimensional reaction coordinate we extracted 28-30 windows in 0.1 nm bins to sample the full 3 nm window. Individual umbrella windows were run for 40 ns under an NVT ensemble with IAPP Cα atoms restrained by harmonic restraints (1000 KJ/mol-nm^2^), resulting in about 55.6 μs of simulation time for PMF calculations. The free-energies over the reaction coordinate were normalized using the Weighted-Histogram Analysis Method (WHAM)^38^ with bootstrapping (N=100) via the GROMACS ‘gmx_wham’ tool.^39^ Binding was then calculated using an in-house python script by taking the potential of mean force difference between the unbound and maximally-bound state, resulting in a Helmholtz free energy. Molecular visualization, coloring, and rendering was subsequently performed using Visual Molecular Dynamics (VMD).^40^

## RESULTS

### Modelling IAPP binder interactions using a simple two-state model

Given the sparsity of experimentally derived structures and their binding sites, we tested IAPP binding independently to the N-terminus (residues 1-22) and C-terminus (residues 19-37) in either a fully α-helical or beta-stranded conformation. For each binder and IAPP target, we ran four technical replicates with different starting coordinates to ensure adequate sampling of the binding event and to overcome any initial bias of the starting system, which sums to 192 binding simulations (48 target-binder complexes x 4 replicates, Figure S1A, S1B). To assess binding progress for each target-binder pair, we monitored the distance between the binder and IAPP backbone (Figure S2). Binding typically converged under 200 ns for IAPP and Aβ foldamers (F_IAPP_ and F_Aβ_) (Figure S2A, Movie 1), although the conformationally restrained β-inhibitors (β_I_), unrestrained β-inhibitors control (β_c_), and PolyA (A_6_), used as controls did not converge in 200 ns (Figure S2B, Movie 2). Instead, these molecules only transiently sampled binding (Figure S2C). Given the computational limitations to sample each binding event over a longer timescale (since 38.4 μs of atomistic MD was used to equilibrate all 192 simulations), we decided to cluster and extract the transient complexes with the highest bound occupancy time (“see *Methods* section for details”). For some simulations, the occupancy times between replicates were heterogenous (Figure S2C, Table 1), which was likely due to the random ligand starting positions in each replicate. Therefore, we decided on a consensus site when the same transient complex was extracted from a majority of the technical replicates. Thus, complexes were extracted from all 48 IAPP-binder permutations either from direct convergence or statistically through the extraction of recurring transient complexes. Taken together, these heuristics (outlined in greater detail in the *Methods* section) allowed us to identify 48 representative complexes containing each binder in the presence of each IAPP target and conformation (Figures S3 and S4).

Next, we ran umbrella sampling (US) simulations on each complex to quantitatively compare the binding affinities between individual ligands and structured IAPP fragments. The resulting potential of mean force (PMF) plots (Figure S5) show the free-energy as the ligand-protein complexes are separated. As expected, the functional form of each PMF resembles a Lennard-Jones potential (Figure S5A) containing both a repulsive and an attractive domain. Since the bound state of the binder was consistently located at the local energy minima, we quantified the free energy of binding (ΔG) for each complex by taking the relative difference in energy between the ligand-free state (d ~ 3 nm) and the ligand-bound state (d = 0 nm) (Figure S5, Figure 2, Table 2). Error bars for each PMF were deduced through bootstrapping, with fluctuations on the order of 1-3 kJ/mol. The IAPP backbone was positionally restrained during pulling, which significantly reduced the time required to sample conformational variation of binders alone, though a combined simulation time of 57.6 μs (40 ns/bin x 30 bins x 48 configurations) was performed overall to deduce the relative affinities of each complex.

**Figure 2.**
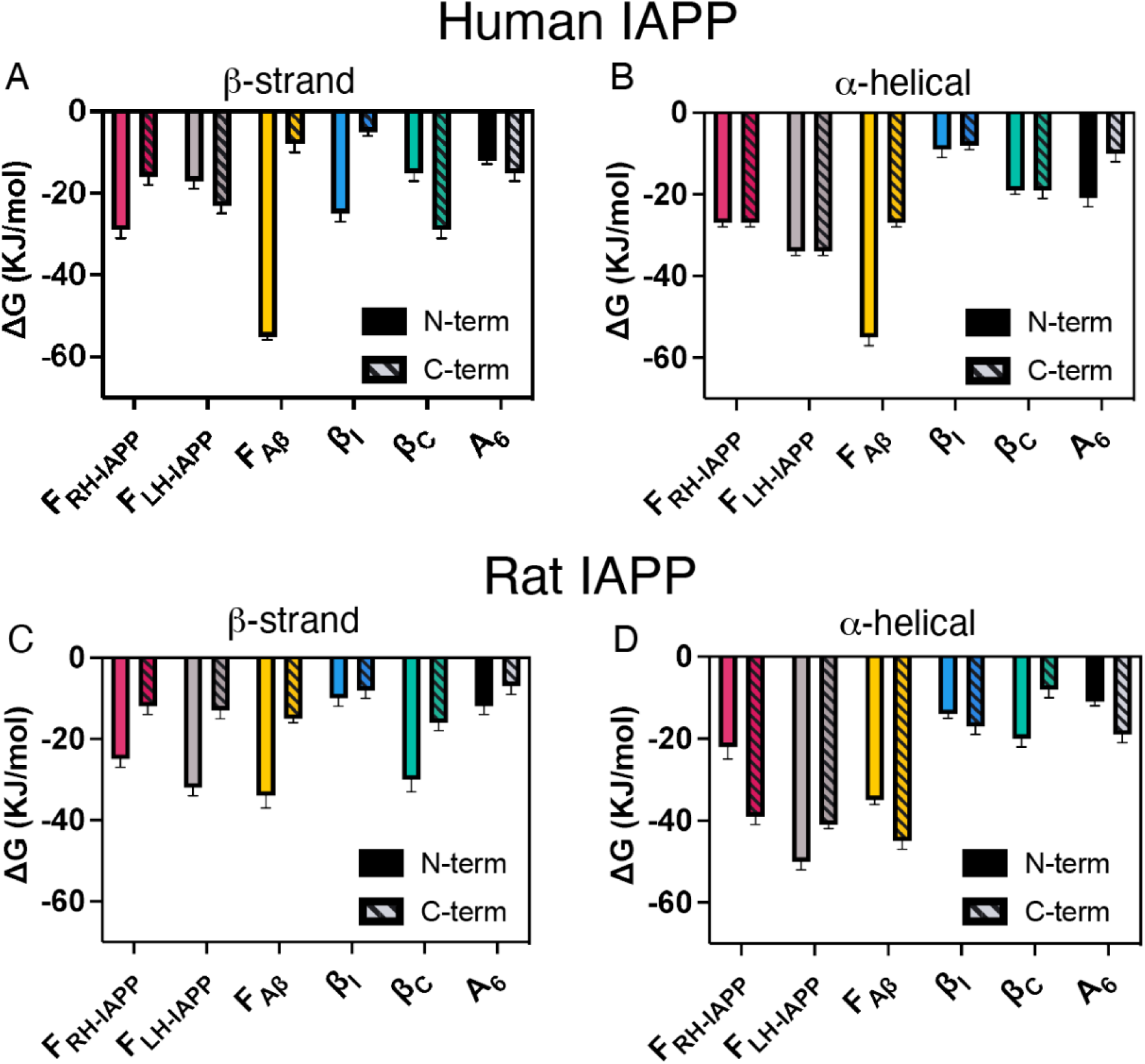
Predicted binding affinities of IAPP binders to N- and C-terminal IAPP from Umbrella Simulations. Shown are inhibitor affinities to **(A, B)** human or **(C, D)** rat IAPP sequences fixed in a **(A, C)** beta-strand or **(B, D)** α-helix conformation. Inhibitors: *right-handed* IAPP Foldamer (F_RH-IAPP_, red), *left-handed* IAPP Foldamer (F_LH-IAPP_, grey), Aβ foldamer (F_Aβ_, yellow), β-inhibitor (β_I_, blue), unmethylated control peptide (β_C_, green), and PolyA (A_6_, black).

### Two-state model is qualitatively consistent with previous experimental data on IAPP foldamer interactions

To assess the performance of our model, we first asked if the observed binding preferences from our model qualitatively agreed with available experimental results in the literature.^17,22^ NMR and circular dichroism (CD) data reported from the Miranker lab suggest that their right-handed (RH) IAPP foldamer (RH-F_IAPP_) binds preferentially to the N-terminus of IAPP and stabilizes helical secondary structure.^22,41^ Consistent with these observations, our model also found RH-F_IAPP_ had a stronger affinity to N-terminal IAPP (27-29 kJ/mol, Figure 2A) compared to C-terminal IAPP (16-27 kJ/mol). Specifically, RH-F_IAPP_ had significantly higher affinity to the helical C-terminus (27 kJ/mol) compared to the β-stranded C-terminus (−16 KJ/mol) (Figure 2B). Thus, our two-state model is consistent with previous experimental observations of RH-F_IAPP_ binding to IAPP.

### Two-state model recapitulates the disruption of aggregation-prone regions by β-breakers

While the exact molecular mechanism of the Gazit lab’s β-breakers^17^ for disrupting IAPP aggregation have not yet been experimentally characterized, they were designed to mimic and disrupt a novel self-aggregating IAPP N-terminal motif (_13_ANFLVH_18_) observed to drive aggregation (therefore are referred to here as β-inhibitors). Therefore, we hypothesized β-inhibitors (β_I_) to exhibit greater affinity for the N-terminus of IAPP, compared to the C-terminus.^17^ Indeed, our models predicted substantially stronger binding affinity of β_I_ for the β-stranded IAPP N-terminus (25 kJ/mol) compared to C-terminus (5 kJ/mol) (Figure 2A, Table 2). Interestingly, β_I_ did not bind the N-terminal motif (Figure 3, *yellow regions*) directly like the conformationally unrestricted control peptide (β_C_, Figure S1; Figure 3C), but instead bound further towards the N-terminus, forming specific hydrogen bonds with THR4 and THR9 (Figure 3A). Together, these results suggest that our model is sufficient to recapitulate the disruption of IAPP self-assembly by the β-breaker.

**Figure 3.**
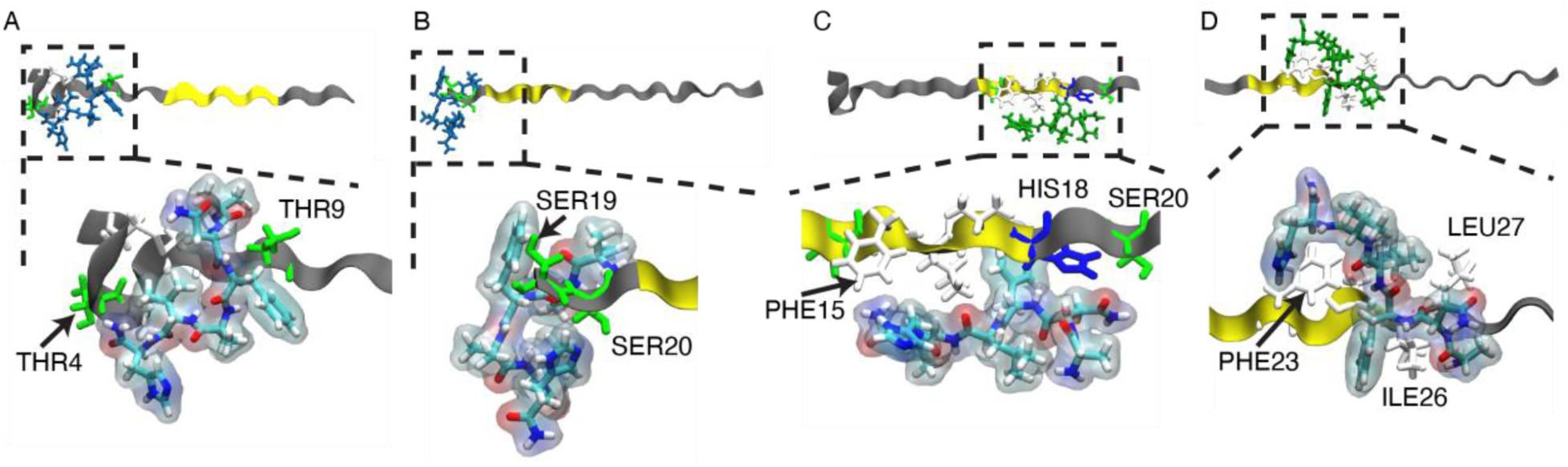
Binding sites of **(A, B)** β-inhibitor and **(C, D)** control peptide to **(A, C)** N-terminal and **(B, D)** C-terminal human IAPP with fixed beta-strand conformation. Aggregation-prone sequences (N-terminal _13_ANFLVH_18_ and C-terminal _22_NFGAIL_28_) on IAPP highlighted in yellow. IAPP residues colored by residue type: hydrophobic (white), polar (green), and positively charged (blue). Foldamer colored by atom type: Carbon (teal), Oxygen (red), Nitrogen (blue), and Hydrogen (white).

### Foldamers interact specifically with IAPP, while poly-arginine control does not

To quantify the degree of specificity and affinity of IAPP binders, we compared them to a hexa-alanine control (A_6_). Alanine is a non-bulky and chemically inert residue, which we hypothesized should make little specific interactions with IAPP (A_6_, Figure S1A). Consistent with this expectation, we found A_6_ binding was generally weak, though it did range between three-orders of magnitude in k_D_ (−7 to −21 KJ/mol; k_D_ of 60-0.2 mM). Given A_6_ primarily interacts only with non-specific backbone and hydrophobic interactions, we expected binders designed for the IAPP sequence to bind with greater affinities. Consistent with our expectations, we find that the IAPP foldamer (F_IAPP_, Figure S1A) had greater affinity for IAPP than A_6_ in all of the complexes in our study. Notably, the F_IAPP_ had an almost three-fold increase in binding affinity for helical C-terminal IAPP (−27 KJ/mol), compared to A_6_ (−10 KJ/mol) (Figure 2B). Additionally, it outperformed A_6_ when binding the beta-stranded N-termini, but not the beta-stranded C-termini of IAPP (Figure 2A).

In our model, A_6_ only transiently interacts with IAPP targets (Figure S2), exhibiting more extended conformations in the presence of beta-stranded IAPP, but more compact states when binding α-helical IAPP (Figures S2, S3). By comparison, RH-F_IAPP_ binds helical IAPP through a combination of hydrophobic interactions between the foldamer oligoquinoline backbone. Specifically, RH-F_IAPP_ interacts with hydrophobic IAPP residues LEU12, PHE15, and LEU16 in the N-terminus (Figure 4A) and TYR37 in the C-terminus (Figure 4A, C). Additional electrostatic interactions between a negatively charged carboxyl group of the foldamer and the positively charged amine of LYS1 help facilitate the binding of RH-F_IAPP_ to the IAPP N-terminus (Figure 4A).

**Figure 4.**
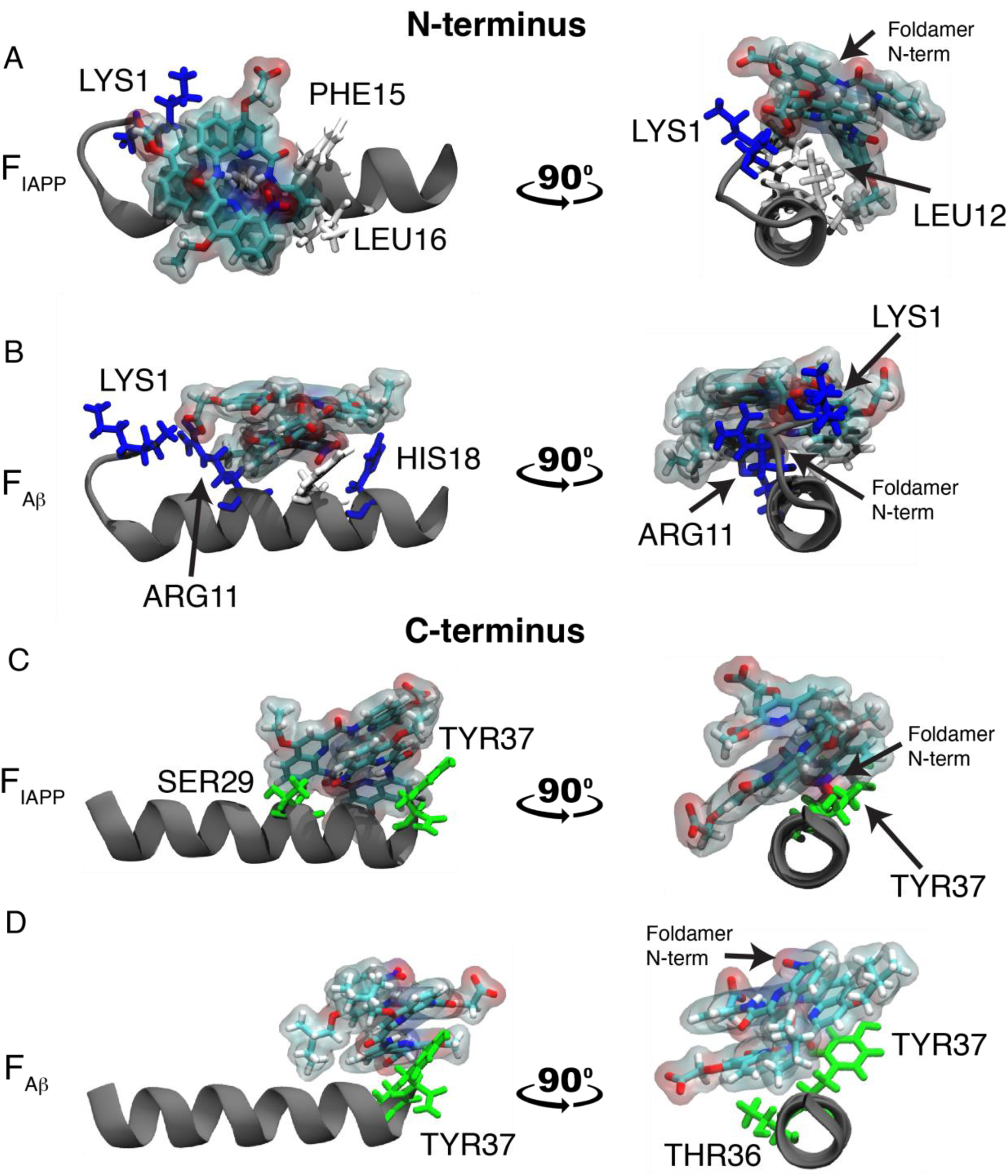
Side-view and helix-view (N-terminus facing out of page) of peptidomimetic foldamers binding **(A, B)** N-terminus and **(C, D)** C-terminus of IAPP after α-helix induction. Inhibitors: **(A, C)** IAPP Foldamer and **(B, D)** Aβ Foldamer. Protein residues colored by type: hydrophobic (white), polar (green), and positively charged (blue). Foldamer colored by atom type: Carbon (teal), Oxygen (red), Nitrogen (blue), and Hydrogen (white).

### Comparing the binding affinity and side-chain interactions of Aβ and IAPP foldamers reveals requirements for reducing IAPP pathology

To better understand the requirements for predicting effective IAPP inhibitors, we compared two foldamers optimized for two different IDP targets (F_IAPP_, which targets IAPP, and F_Aβ_, which targets amyloid beta, Figure S1A). While F_IAPP_ differs from F_Aβ_ only by the conservative replacement of two ethyl groups with slightly larger 2-methyl butane groups (Figure S1A), only F_IAPP_ rescues cells from IAPP-induced cell toxicity^15^. We hypothesized that the more sterically demanding side chains of F_Aβ_ may disrupt binding to IAPP. Surprisingly, our model predicted F_Aβ_ to bind the IAPP N-terminus with significantly more affinity (55 kJ/mol or ~0.3 nM) compared to F_IAPP_ (28 kJ/mol or ~13 μM) in both β-stranded and alpha-helical conformations (Figure 2A, B). For C-terminal IAPP, F_Aβ_ exhibited only half of the binding affinity to beta-stranded IAPP (−8 KJ/mol) compared to F_IAPP_ (−16 KJ/mol) and equal affinity to alpha-helical IAPP (27 kJ/mol or ~300 μM) (Figure 2A, B). Further, the larger steric requirement of the F_Aβ_ sidechains is well accommodated and only slightly modifies the most stably observed binding site (Figure 4A-D). The increase in binding affinity of F_Aβ_ to the IAPP N-terminus compared to F_IAPP_ seems to be partly driven by the additional hydrophobicity of the foldamer, as indicated by a significant increase in bound surface area (BSA) (Table 3). Additionally, we observed a bias for F_Aβ_ to bind IAPP in the opposite orientation (via C-terminal methyl-ester) than F_IAPP_ (via N-terminal nitro-group), which exposes the more polar end of the foldamer to establishing hydrogen bonds with the positively charged IAPP residues LYS1, ARG11, and HIS18 (Figure 4A, B). Taken together, our model reveals differences in the selectivity of binding between two chemically similar foldamers and highlights the plausible mechanisms by which they differentially interact with their targets.

### Foldamers can bind rat orthologs, but not beta-inhibitors or PolyA

To test the extent to which mutations disrupt the ability of the hIAPP binders to bind pre-induced conformations, we formed analogous complexes with rat IAPP (rIAPP) and measured the binding affinity between each binder and rIAPP peptides using the same protocol as before. As expected, the range of binding affinities of our poly-alanine control (A_6_) for rIAPP (−7 to −19 KJ/mol) did not significantly change compared to human IAPP in either conformation (−7 to −21 KJ/mol), consistent with its ability to form only non-specific interactions with the peptide targets (Figure 2C, D, Figure S4). By comparison, the right-handed IAPP foldamer (RH-F_IAPP_) exhibited much stronger affinities for rIAPP (−12 to −39 KJ/mol) than A_6_ (Figure 2C, D; Table 2). Consistent with model’s previous predictions that RH-F_IAPP_ does not interact with HIS18 in either conformation of the hIAPP N-terminus (Figure 4, Figure S3), the binding of RH-F_IAPP_ to rIAPP N-termini was similar to hIAPP in either β-stranded or helical conformations (Figure 2C, D). However, in the C-terminus, RH-F_IAPP_ binding was two-fold greater (−39 KJ/mol; ~160 nM) than A_6_ (−19 KJ/mol), and greater than its binding for hIAPP C-terminus (−27 KJ/mol). We also observed similar ranges of binding affinities of the amyloid-beta targeting foldamer (F_Aβ_) to rIAPP (−15 to −45 KJ/mol), compared with its binding to hIAPP (−8 to −55 KJ/mol) (Figure 2C, D; Table 2). Therefore, we find foldamers largely retain binding for pre-induced IAPP conformations, despite sequence differences between hIAPP and rIAPP.

Given the similarity of the β-inhibitor (β_I_) sequence to the self-assembling region of IAPP (_13_ANFLVH_18_), we hypothesized the H18R mutations in the N-terminus could be disruptive to its binding. Indeed, our model predicted a decrease in binding of β_I_ to the β-stranded rIAPP N-terminus (−10 KJ/mol), which was much stronger for hIAPP (−25 KJ/mol). The binding affinities of β_I_ to the rest of the rIAPP targets were similar to A_6_, which was also observed for their binding to hIAPP (Figure 2C, D; Table 2). Consistent with our expectations, the H18R mutations disrupted the ability of the conformationally unrestricted peptide control (β_C_) from associating with residues 13-18 as it did with hIAPP (Figure 3C; Figure S3; Figure S4). Interestingly, the model predicts the new binding site further towards the rIAPP N-terminus has double the binding affinity for the β-stranded rIAPP N-terminus (−30 KJ/mol) than for hIAPP (−15 KJ/mol) (Figure 2A, C; Figures S3, S4). Given that this sequence was not modified in rIAPP and is also present in the hIAPP, we asked why it was not sampled in the hIAPP simulations. One plausible explanation is transient binders like β_I_ and β_C_ exhibit significantly lesser bound occupancy times (10s of ns) than the foldamers (up to 100s ns), and thus, sample less IAPP surface area than the foldamers during the same simulation time (Table 1). Despite these limitations, our model clearly demonstrates that binding of β_I_ is disrupted by the rat mutations, suggesting their binding requires the presence of specific hIAPP sequence motifs.

### IAPP Foldamer handedness does not affect conformational selection but changes its sequence selection

Previous CD and Isothermal Calorimetry (ITC) experiments revealed F_IAPP_ can interconvert between a right-handed (RH) and left-handed (LH) conformation, yet only RH-F_IAPP_ binds strongly to hIAPP.^22^ Therefore, we asked if our model could distinguish between the affinities of the two conformers and help explain this interesting binding behavior. To answer this question, we measured the binding affinity of both foldamer conformers to the same IAPP targets. We first noted F_IAPP_ conformer interconversion did not occur during the timescale of our simulations (Figure S6). This is notable because isolating RH-F_IAPP_ and LH-F_IAPP_ experimentally requires the addition of chiral groups to prevent the interconversion of F_IAPP_ conformations^22^. Therefore, our model can test the binding of individual F_IAPP_ conformations without these chemical modifications. For beta-stranded IAPP, we observed a significant reduction of binding of LH-F_IAPP_ (−17 KJ/mol) to N-terminal IAPP compared to RH-F_IAPP_ (−29 KJ/mol) (Figure 2A). For the C-terminus, there was a moderate increase in affinity of LH-F_IAPP_ compared to RH-F_IAPP_, revealing conversion of RH-F_IAPP_ to LH-F_IAPP_ can switch sequence selectivity from the beta-stranded hIAPP N-terminus to C-terminus (Figure 2A). This conformer selectivity was not observed for rIAPP, where the RH- and LH-F_IAPP_ had similar preferences (Figure 2C).

For helical IAPP, the model predicted similar selectivity between RH- and LH-F_IAPP_ binding, with LH-F_IAPP_ exhibiting moderately higher affinity for both hIAPP termini and significantly higher affinity for binding of LH-F_IAPP_ to helical N-terminal rIAPP (50 KJ/mol) compared to RH-F_IAPP_ (−22 KJ/mol) (Figure 2B, D). This increase in affinity of LH-F_IAPP_ for helical N-terminal was attributed to its ability to interact find a stable interaction with all three positively charged N-terminal IAPP residues (LYS1, ARG11, and HIS18), whereas RH-F_IAPP_ only strongly interacts with LYS1. Taken together, our model reveals both RH- and LH-foldamer conformers strongly bind helical IAPP as well as some beta-stranded targets, suggesting the preference of foldamer handedness during IAPP binding is not driven by conformational selection alone, but may require induced-fit binding modes beyond the scope of our model.

### Binding of IAPP Foldamer is conformationally selective, but β-breaker is driven by both sequence and conformational selection

Lastly, given the ability of our model to measure *conformation-specific* binding affinities, we wanted to calculate the conformational selection (defined as the change in binding between specific conformations for a fixed sequence) and sequence selection (defined as the change in binding between specific sequences for a fixed conformation) of each binder. By decoupling these normally experimentally indistinguishable processes, we can predict which dominates the binding behavior we observe. We hypothesized that the molecules with previously observed experimental efficacy to reduce hIAPP aggregation will have greater conformational selection to hIAPP than controls that do not. To test this hypothesis, we calculated both the conformational selection (ΔΔG_CONF=_ ΔG_⍺_ − ΔG_β_) and sequence selection (ΔΔG_SEQ_ = ΔG_C_ – ΔG_N_) of each complex (where ΔG_⍺_ and ΔG_β_ are the binding affinities to ⍺-helical and beta-stranded conformations, and ΔG_C_ and ΔG_N_ are the binding affinities to the hIAPP N- and C-termini, respectively) (Figure 5A). Using these four values, we can classify binders as conformationally selective if the absolute value of ΔΔG_CONF_ is large and the absolute value of ΔΔG_SEQ_ is small. Alternatively, when the absolute value of ΔΔG_CONF_ is small and the absolute value of ΔΔG_SEQ_ is large, the binder is classified as sequence selective. Further, we distinguish between *indiscriminate* and *sequence-conditional* binders by the ability to maintain its preference for a conformation despite changes in sequences (e.g. N-terminus vs. C-terminus, hIAPP vs. rIAPP). It is also important to note that each ΔG is negative/adhesive (as in Table 2), consistent with the nomenclature above.

**Figure 5.**
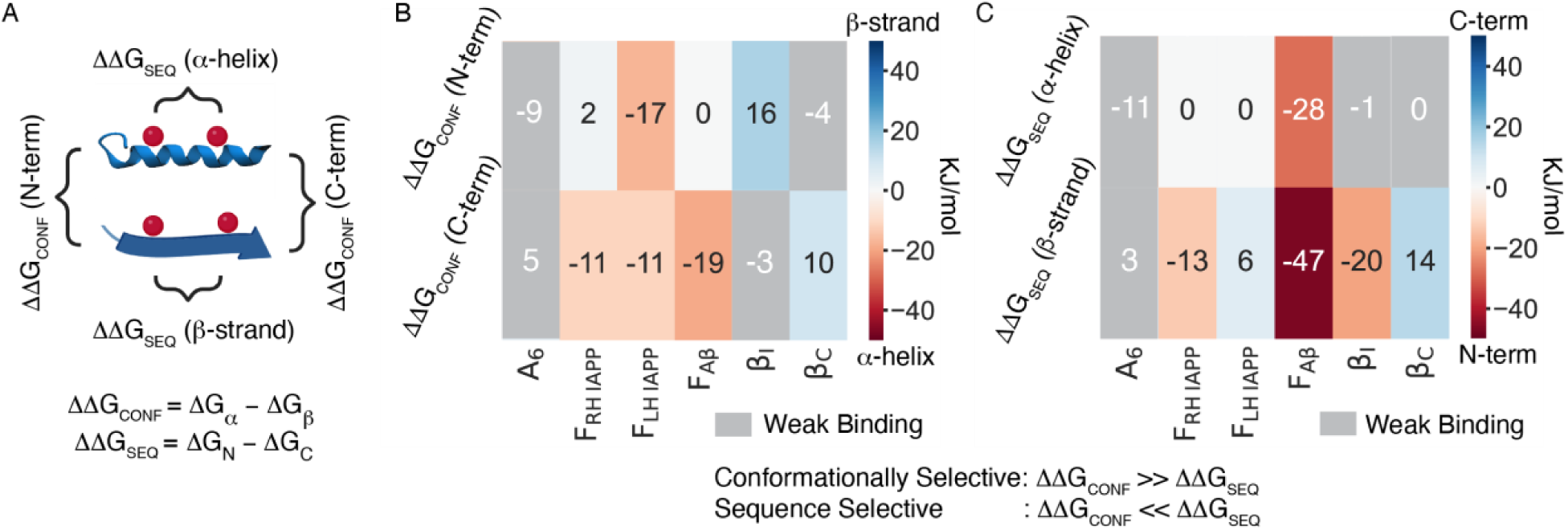
**(A)** Conformation and sequence preferences for each binder calculated from differences in ΔG of binding (ΔΔG) to *human* IAPP. **(B)** Conformational preference of IAPP binders, where α-helical binders are red and β-sheet binders are blue. **(C)** Sequence preference of IAPP binders, where N-terminal binders are in red and C-terminal binders in blue. A_6_ and binders with less binding affinity than A_6_ (“weak binders”) are grayed out.

The colormap in Figure 5 displays the ΔΔG_CONF_ and ΔΔG_SEQ_ for all molecules tested in our simulations. Binders that did not stably bind IAPP, such as the polyA control (A_6_), or were below the binding affinity of A_6_ (Figure 2) were not considered and are shown in grey (Figure 5B, C). Our first observation was that the binders showed different degrees of conformational selection depending on the target sequence, suggesting conformational selection may be coupled to the target’s sequence. Interestingly, some binders maintained their conformational selection despite differences in sequence between the N- and C-termini, while others lost their conformational selectivity (Figure 5B, 5C). We classified left-handed IAPP foldamer (LH-F_IAPP_) as an *indiscriminate* conformationally selective binder, because it has conformational selection in both the N-terminus (−17 KJ/mol, Figure 5B, *top*) and C-terminus (−11 KJ/mol, Figure 5B, *bottom*) of IAPP. In contrast, the right-handed IAPP-foldamer (RH-F_IAPP_) and the amyloid-beta foldamer (F_Aβ_) are binders with *sequence-conditional* conformational selection, exhibited only for the C-terminus (Figure 5B). Similarly, the β-inhibitor (β_I_) and peptide control (β_C_) also have conformational selection only in one terminus, suggesting they are also *sequence-conditional* conformationally selective binders.

Another consideration is the relative magnitudes of ΔΔG_CONF_ and ΔΔG_SEQ_, which determine if binders are driven by conformation or sequence selection. We compare the ΔΔG_CONF_ of binders to the appropriate ΔΔG_SEQ_, depending on what conformation is preferred. For example, LH-F_IAPP_ has conformational selection for helical hIAPP at both termini (−11 and −17 KJ/mol, Figure 5B). Therefore, we compare these values to its ΔΔG_SEQ_, to the value of ΔΔG_CONF_ for hIAPP induced in a helical conformation (0 KJ/mol, Figure 5C, *top*), which reveals its binding is largely driven by conformational selection. For conditional binders like RH-F_IAPP_, which are only conformationally selective for helical hIAPP in the C-terminus, we compare its ΔΔG_CONF_ (−11 KJ/mol, Figure 5B, *bottom*) in the C-terminus to its ΔΔG_SEQ_ (0 KJ/mol, Figure 5C, *top*) for the helical conformation, again revealing that its binding is driven by conformational selection. In contrast, conformational selection of the amyloid-beta foldamer (F_Aβ_) for helical hIAPP was much smaller (−19 KJ/mol, Figure 5B, *bottom*) than its ΔΔG_SEQ_ (−28 KJ/mol, Figure 5C, *top*) for the helical conformation, suggesting its sequence selection may overshadow its conformational preferences.

Lastly, the binding of β_I_ and β_C_ is a result of both ΔΔG_CONF_ and ΔΔG_SEQ_. β_I_ exhibits high ΔΔG_CONF_ (−16 KJ/mol, Figure 5B, *top*) for beta-stranded hIAPP and also high ΔΔG_SEQ_ (−20 KJ/mol, Figure 5B, *bottom*) for this fixed conformation. Similarly, β_C_ exhibits high ΔΔG_CONF_ (10 KJ/mol, Figure 5C, *bottom*) for beta-stranded conformations, and high ΔΔG_SEQ_ (14 KJ/mol, Figure 5C, *bottom*) for this fixed conformation. Thus, both conformation and sequence selection are involved in binding of β_I_ and β_C_ to hIAPP, with slight preference for sequence selectivity. Given the poor affinity of β_I_ for C-terminal and helical IAPP targets (Figure 2), we did not draw comparisons between ΔΔG_CONF_ and ΔΔG_SEQ_ (Figure 5B, C, *gray*). Similarly, we did not draw comparisons between ΔΔG_CONF_ and ΔΔG_SEQ_ due to the poor binding affinity of β_C_ to N-terminal and helical IAPP targets (Figure 2) (Figure 5B, C, *gray*). Taken together, β-breaker peptide binding to hIAPP is driven by an equal contribution of both sequence and conformational selection. Quantitatively analogous trends of all binders in this study can be found for rIAPP in Figure S7. A summary of the selectivity of all binders in this study to hIAPP and rIAPP is shown in Table 4.

## DISCUSSION

Islet amyloid polypeptide (IAPP) is an example of an intrinsically disordered proteins (IDP), a class of proteins which make up approximately a third of the human proteome and are involved in essential biological processes such as transcription, translation, and cellular signalling.^42^ The close relationship between IDPs and human diseases make them attractive drug targets, however; they were thought to be too dynamic and lack the defined binding pockets necessary for small-molecule binding.^2^ The recent design of folded small molecules (i.e. foldamers) demonstrated at least one way to modulate IDP ensembles, by targeting transient helical intermediates preventing pathological misfolding.^41^ Using the foldamers, in part, as inspiration, we designed this study to more broadly assess the feasibility of drugging IDPs by optimizing molecules for a specific conformation.

In this work, we demonstrate our two-state model can reliably predict the binding preferences of two classes of IAPP-targeting molecules for a particular conformation. The foldamers used in this study were designed to target helical hIAPP intermediates and shown to stabilize these structures derived from NMR and CD data.^22,41^ Consistent with these previous experimental results, our model predicts that foldamers have strong binding to α-helical conformations in both IAPP termini (Figure 2). Further, β-breakers (referred to as β-inhibitors, β_I_, in this study) for IAPP were designed to bind and disrupt the self-interactions of sequences known to form beta-sheet structure and thus prevent the assembly of amyloid oligomers and fibrils.^17^ Our models captured this preference for the β-inhibitors as it exclusively bound the hIAPP N-terminus, though in a region different from the predicted nucleation region (Figure 2, Figure 3). Therefore, our models faithfully recapitulate biophysical observations from previous experiments while providing molecular insight into their mechanisms and dynamics of binding.

The binding of the foldamers appears to be driven by a combination of conformational selection and induced-fit, while the β-inhibitors are driven by primarily conformational selection. Specifically, foldamers have conformational selection for ⍺-helical conformations in the C-terminus, but not in the N-terminus (Figure 2). This suggests that stabilization of the N-terminal helix may require adaptive fitting, consistent with an induced-fit mechanism where initial binding is further stabilized by conformational rearrangements. This conformational rearrangement could then enhance binding of the foldamer to the C-terminus of IAPP, which was usually much weaker than binding to the N-terminus (Figure 2B, D). In contrast, the preference for beta-stranded IAPP of the β-inhibitor can be explained by conformational selection alone (Figure 5). These findings explain the functional diversity between these two-classes of inhibitors. While the function of the β-inhibitor has not been tested *in vivo*, its application reduces the number of aggregates without fully preventing aggregation. In contrast, the foldamer has been shown to completely inhibit the formation of fibrils for at least 40h, reduce IAPP-induced membrane leakage, and protect against mitochondrial depolarization.^15^

Another difference between the IAPP-targeting foldamer and β-inhibitor lies in their sequence selectivity; the foldamers bind both IAPP termini while the β-inhibitor only binds the N-terminus (Figure 2). Based on the aforementioned experimental data and our simulations, we conclude that binding to both IAPP termini is required to reduce its pathological effects (Figure 2, 4, 5). Further evidence of this hypothesis is provided from the design of interaction surface mimics (ISMs) such as IAPP-GI, a full-length IAPP mimic that has both N- and C-terminal aggregation-prone regions.^43^ Notably, one of these regions contains two N-methylated residues to block aggregation and reduce cytotoxicity.^44^ Interestingly, both the ISMs and foldamers can stabilize dynamic or liquid-like oligomers, instead of off-pathway fibrils or discrete complexes.^16,44^ Therefore, it seems plausible that the ability of foldamers and ISMs to promote inter-molecular interactions with the inhibitor, rather than intra-molecular interactions, stabilizes the IAPP monomer in an expanded conformation and prevents it from accessing pathological conformations.

Foldamers are predicted to be more promiscuous binders than β-inhibitor peptides. Surprisingly, foldamers had strong binding to both hIAPP and the non-aggregating rat IAPP (rIAPP) control peptides, especially for ⍺-helical conformations (Figure 2B, D). By comparison, the β-inhibitor’s ability to bind the hIAPP N-terminus in a beta-stranded conformation was lost (Figure 2A, C). This is interesting because it suggests the foldamers are less sensitive to the rIAPP mutations than the β-inhibitor. This observation might be due to differences in their rational design. The foldamers were designed to putatively interact with complementarily charged residues in an ⍺-helical conformation, which were previously shown to be formed between residues 9-22 in the presence of anionic lipids.^41,45^ Since only the H18R mutation is within this range, it was not surprising that the foldamer had similar affinities between human and rat IAPP (Figure 2). We did not test a non-IAPP target sequence due to computational limitations, but this would be interesting to test in the future to assess the foldamer’s propensity to bind off-target sequences with the identical conformation.

This enhanced affinity of F_Aβ_ for the IAPP N-terminus may stem from having more hydrophobic area to contribute to binding (Figure S1), evident by its greater bound surface area compared to the F_IAPP_ (Table 3). Further, the similarity of the Aβ peptide, which shares 50% sequence similarity with IAPP, may contain sequence-encoded recognition motifs both foldamers recognize.^47,35^ This implies that foldamers may exhibit cross-interactions with similar IDPs, potentially enabling simultaneous targeting of multiple misfolding proteins. Taken together, our results reveal how tuning of the foldamer sidechains are critical for optimizing conformational selection and generating binders that prevent IAPP toxicity.

In contrast to the foldamers, our models and previous experiments^17,46^ reveal the disruption of the β-inhibitor binding to rIAPP is three-fold. Firstly, the conformational restriction prevents expanded beta-stranded templating to IAPP and in most cases lower affinity (Figure 2, S3, Table 2). Secondly, the H18R mutation in rIAPP disrupts interactions with HIS18, as observed by the non-conformationally restricted control peptide (β_C_, Figure 3C). Lastly, rIAPP does not natively adopt beta-stranded conformations, thought to be a result of proline mutations in the C-terminus.^46^ Our models of the β-inhibitors reveal that even when rIAPP is fixed in the beta-stranded conformation, the affinity is quite lower (Figure 2A, C). Taken together, we provide molecular insights into the loss of β-inhibitor binding to rIAPP and demonstrate they have greater sensitivity to rIAPP mutations than the foldamers.

In addition to optimization binders based on their binding affinity to a target structure, it must also have conformational selection—the ability to distinguish that conformation from others. We found that optimizing binders based on binding affinity alone is not sufficient to predict the rescue of IAPP toxicity. If we were to select a binder based on affinity alone (Figure 2), our results would suggest the non-cytoprotective Aβ-foldamer (F_Aβ_) to be the ‘best’ binder, which had stronger binding than the IAPP Foldamer (F_IAPP_) to almost every IAPP target we tested (Figures 2, 5). However, if we consider the relative contributions of conformation-(ΔΔG_CONF_) and sequence-selection (ΔΔG_SEQ_), F_IAPP_ stands out as having similar magnitudes of ΔΔG_CONF_ and ΔΔG_SEQ_. By comparison, the F_Aβ_ binding is primarily dominated by ΔΔG_SEQ_, thus sequence selection overshadows its conformational preferences (Figure 5). Therefore, only the IAPP Foldamer is optimized for hIAPP, where both sequence selection and conformational selection are equal determinants in binding.

For the β-inhibitor, conformational selection is mediated by specific side-chain interactions (Figure 3). Backbone interactions alone (e.g. A_6_) did not impart conformational selection nor bind effectively (Figures 2, 5). Interestingly, the sequence specificity of the control peptide (β_C_) for the C-terminus can be switched to a non-canonical N-terminus motif by the addition of the methyl groups in the β-inhibitor (Figure 3). This effect can be attributed to the conformational restriction imposed by the β-inhibitor that prevents forming parallel β-sheet interactions with known IAPP nucleation sites (Figure 2, Figure 3). Furthermore, the conformational restriction of the β-inhibitor likely reduces templating of the IAPP N-terminus, which has been shown to promote the misfolding of disordered monomers on folded scaffolds, such as amyloid fibrils.^47^ The methylation of the peptide may also act as a steric inhibitor, similar to the Eisenburg’s N-methylated IAPP inhibitors based on a Cryo-EM map of the fibril.^20^ The limitation to these class of inhibitors is that they have a lower binding affinity than their non-methylated counterpart^20^, a phenomenon also captured in our model (Figure 2). However, our work indicates that methylated inhibitors such as the ones examined, can bind with conformational selection, thereby targeting only misfolded, non-functional IAPP.

Future models should consider not only the selection of the inhibitor for an IDP, but also how IDPs respond to foldamer binding (induced-fit). While our work demonstrates clear advantages of using fixed-conformational models to measure conformational selection of the inhibitors, it also reveals limitations. Specifically, the inability of the model to distinguish between the handedness of foldamers and explain the lack of foldamer binding to rIAPP (Figure 2), both observed experimentally.^22^ These discrepancies from experiments are likely a result of using fixed-conformation models and highlight the need to incorporate induced-fit binding mechanisms into future models directly.

Additionally, two critical assumptions underpin our model: a) the conformational selection of the inhibitor is driven by differential binding to secondary structure, and b) all residues are equally exposed, assuming a fully extended conformation. Although small IDPs such as monomeric IAPP largely meet these assumptions, it has been demonstrated that IAPP can shield its hydrophobic regions from solvent accessibility when it forms structures such as a complex β-hairpin or in heptameric ⍺-helical micelles.^48,49^ These configurations may facilitate membrane poration.^16,48,50^ Furthermore, larger IDPs can form multivalent weak intra- and inter-molecular interactions, primarily held together by aromatic or electrostatic forces.^8,51^ These interactions can lead to the formation of tertiary structures that potentially inhibit the binding of small molecule inhibitors.

Experimentally validating these predictions is challenging due to the inherent instability of any single IDP conformation. However, we can measure how optimizing around fixed-conformations improves binding to the IDP ensemble by employing standard binding assays such as isothermal titration calorimetry (ITC).^52^ This technique allows us to quantify the binding affinity and thermodynamics of IDP binders, providing insights into how well our model predictions align with actual binding behaviors. Additionally, NMR and CD experiments can verify the predicted stabilization of IDP secondary structures, as has been conducted in previous studies.^22,35^

To validate the conformational selection of binders, we may be able to use kinetic methods, such as stopped-flow,^53^ and single-molecule fluorescent approaches^54^ to monitor the structural transitions of IDPs and their co-assemblies upon binding, offering detailed insights into the dynamic behavior of these interactions at the single-molecule level. These approaches provide a powerful means of observing the often transient and complex interactions characteristic of IDP.^54^ Beyond drug design, this work also contributes to the development of environmentally sensitive IDP biosensors.^55^ For instance, sensors designed to be activated only in the presence of specific stabilizers, could be invaluable for identifying how IDPs are biologically regulated. Such tools could also serve to validate the stabilizing capabilities of IDP binders within cellular environments. By combining these experimental methods with molecular models, we can enhance our understanding of the molecular mechanisms of IDP binders, thereby opening new avenues for both therapeutic and diagnostic applications.

## CONCLUSIONS

In summary, this work highlights the role of conformational selection towards the design of binders that can reduce IAPP pathology. We demonstrated that binders optimized for a specific IAPP conformation effectively stabilize that conformation. To illustrate this, we built two models of IAPP in either a ⍺-helical or β-sheet conformation, and measured the binding affinities of known IAPP-targeting inhibitors and controls. Our findings confirm that the model could accurately predict the conformational selection of both classes of molecules tested, validating our hypothesis that conformational stabilization can be predicted by measuring conformation-specific affinities through MD simulations. By leveraging the molecular resolution of our model, we could distinguish between the mechanisms of the inhibitors by revealing crucial interactions that drive their association with each conformation. Lastly, by combining previous experimental data on the functional activity of IAPP inhibitors with our own measurements of conformation- and sequence-specific affinities (ΔΔGs), we demonstrate the ability to predict whether binders are conformation or sequence selective, and to identify the properties that an IAPP binder should possess to maximize its impact on IAPP function. Our findings indicate that, alongside sequence specificity, conformational selection is required for IAPP cytoprotection. Taken together, this work reveals how conformational selection can lead to the stabilization and functional modulation of an IDP, such as the prevention of diabetes-associated IAPP misfolding that leads to β-cell toxicity. We believe our approach has the potential to enable the prediction of novel conformationally selective probes that can distinguish between functional and pathological conformations of IDPs, or even target them for specific degradation.

## Author contributions

B.A.B. and Z.A.L. conceived the study. B.A.B. conducted all GROMACS simulations and analysis. B.A.B. and Z.A.L. wrote the manuscript. Z.A.L. supervised the study.

## Materials and correspondence

N/A

## ACKNOWLEDGMENTS

The authors acknowledge critical discussion with members of the Levine Lab at Altos Labs and Dr. Andrew Miranker at Yale University regarding the manuscript. The authors thank the Computational Research Support Group at Yale’s Center for Research Computing and the Altos Labs Scientific Applications and Computing Team for their technical support. B.A.B. acknowledges support by the Yale MB&B Biophysics Training Grant.

## SUPPLEMENTAL INFO (in EXCEL file)

**Attachment 1.** HBonds, Bound Occupancy Times, Residue, and Convergence Criteria for Binding Simulations.

