## Supplemental Figures for "Drugging Disordered Proteins by Conformational Selection to Inform Therapeutic Intervention"

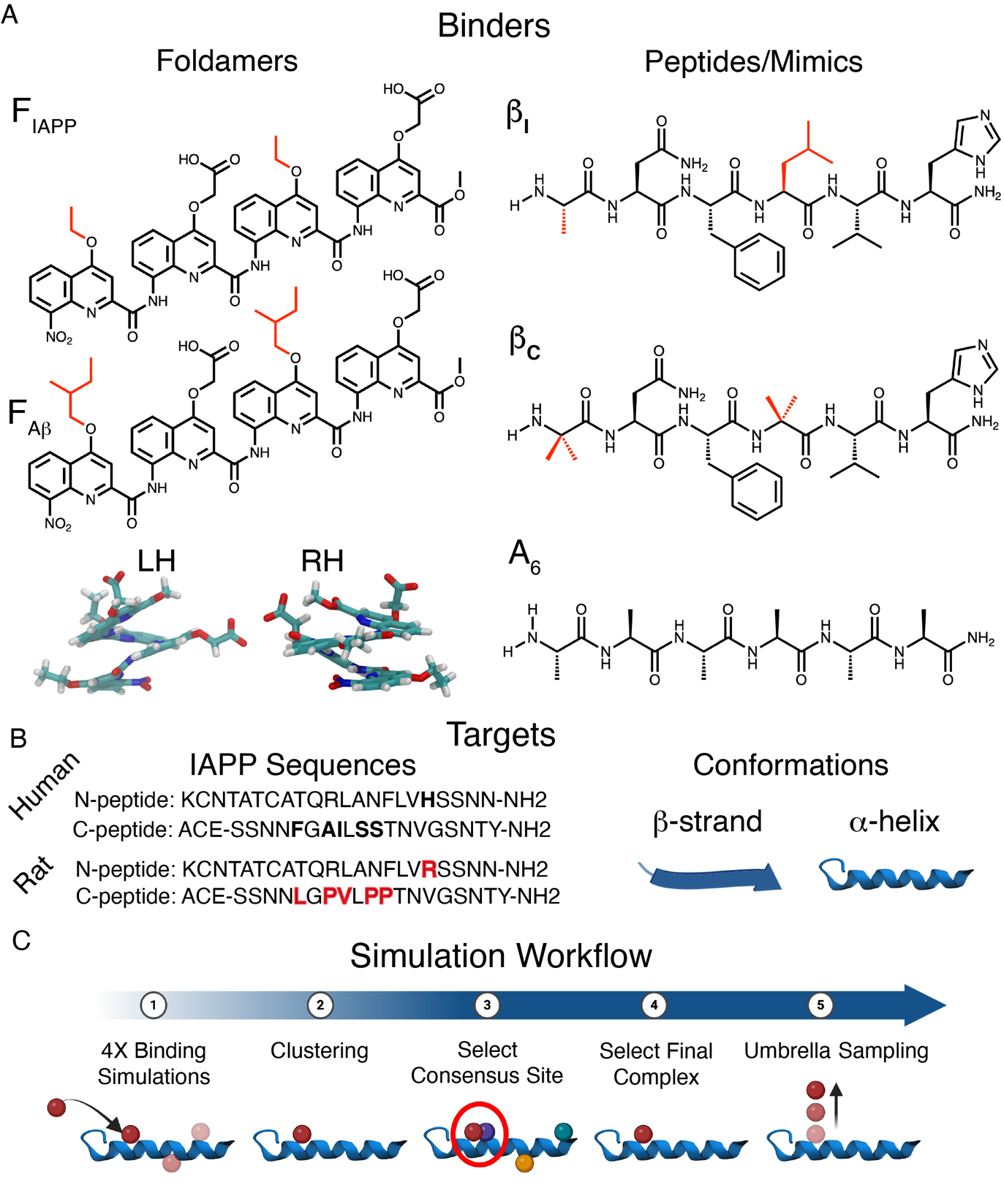
**Figure S1. (A)** Classes of binders tested in this study: IAPP-Foldamer (F_IAPP_), Aβ-Foldamer (F_Aβ_), β-inhibitor (β_I_); and control peptide (β_C_). Modified chemical groups are labelled in red. **(B)** IAPP sequences and conformations used to test binder conformation and sequence selection. Differences between human (black) and rat (red) IAPP sequence are bolded. **(C)** Computational workflow for binding simulations and clustering to extracting final complexes. Binders were each subject to four independent binding simulations. Clustering was used to select the consensus site. The stability of the final complexes was evaluated with Umbrella Sampling.

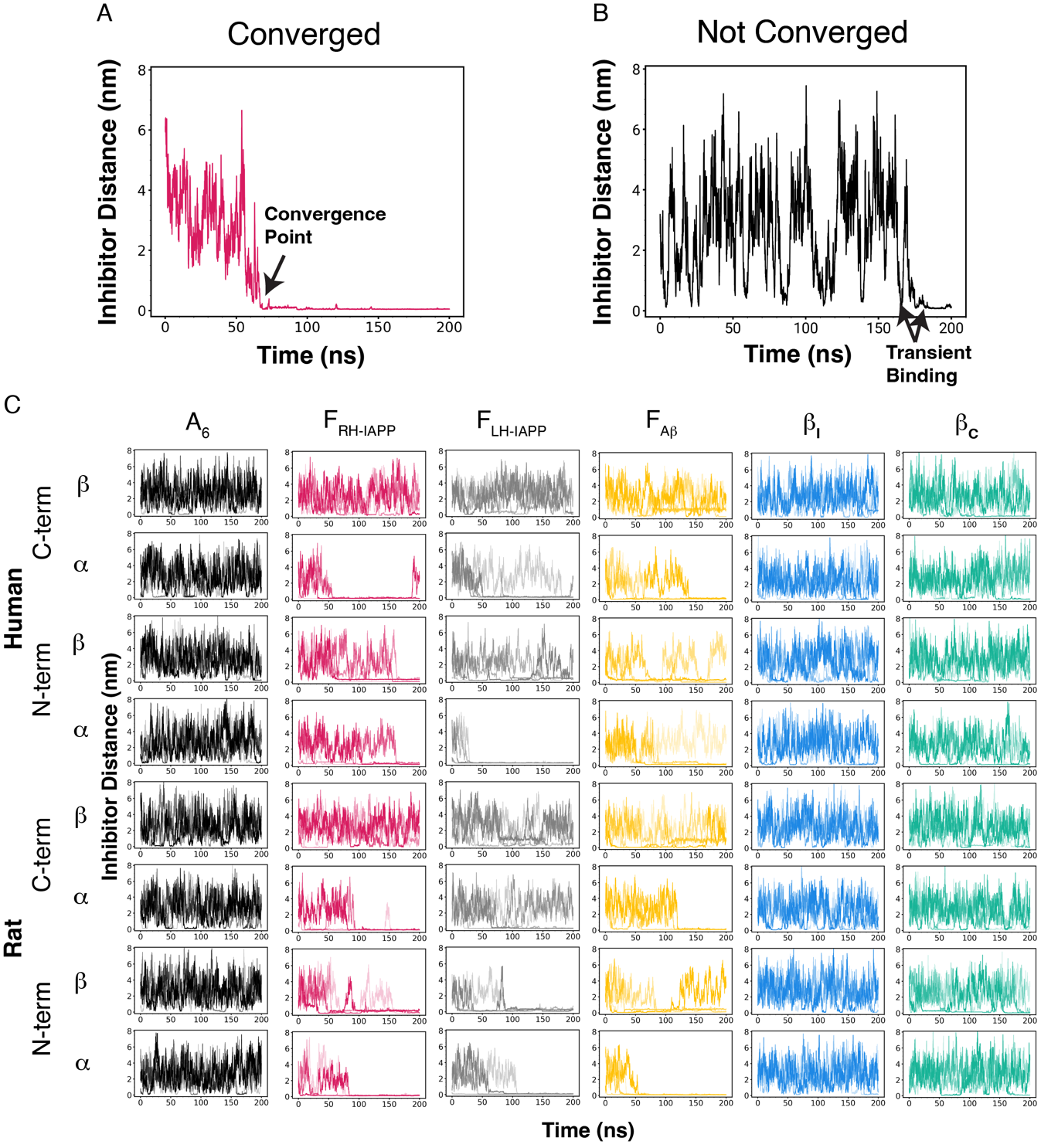

**Figure S2.** Distance of the inhibitor from IAPP backbone over 200ns binding simulation. **(A)** Representative converged simulation, where binder (F_IAPP_) reaches convergence around 60ns. **(B)** Representative non-converged simulation, where binder (PolyA) transiently binds and unbinds IAPP. **(C)** Inhibitor distances calculated over 200ns for all simulations in this study. Each inhibitor column is distinguished by color: PolyA (A_6_, black); RH-IAPP Foldamer (F_RH-IAPP_, red); LH-IAPP Foldamer (F_LH-IAPP_, grey); Aβ-Foldamer (F_Aβ_, yellow); β-inhibitor (β_I_, blue); and control peptide (β_C_, green). Each row represents a different target sequence with a given fixed secondary structure.

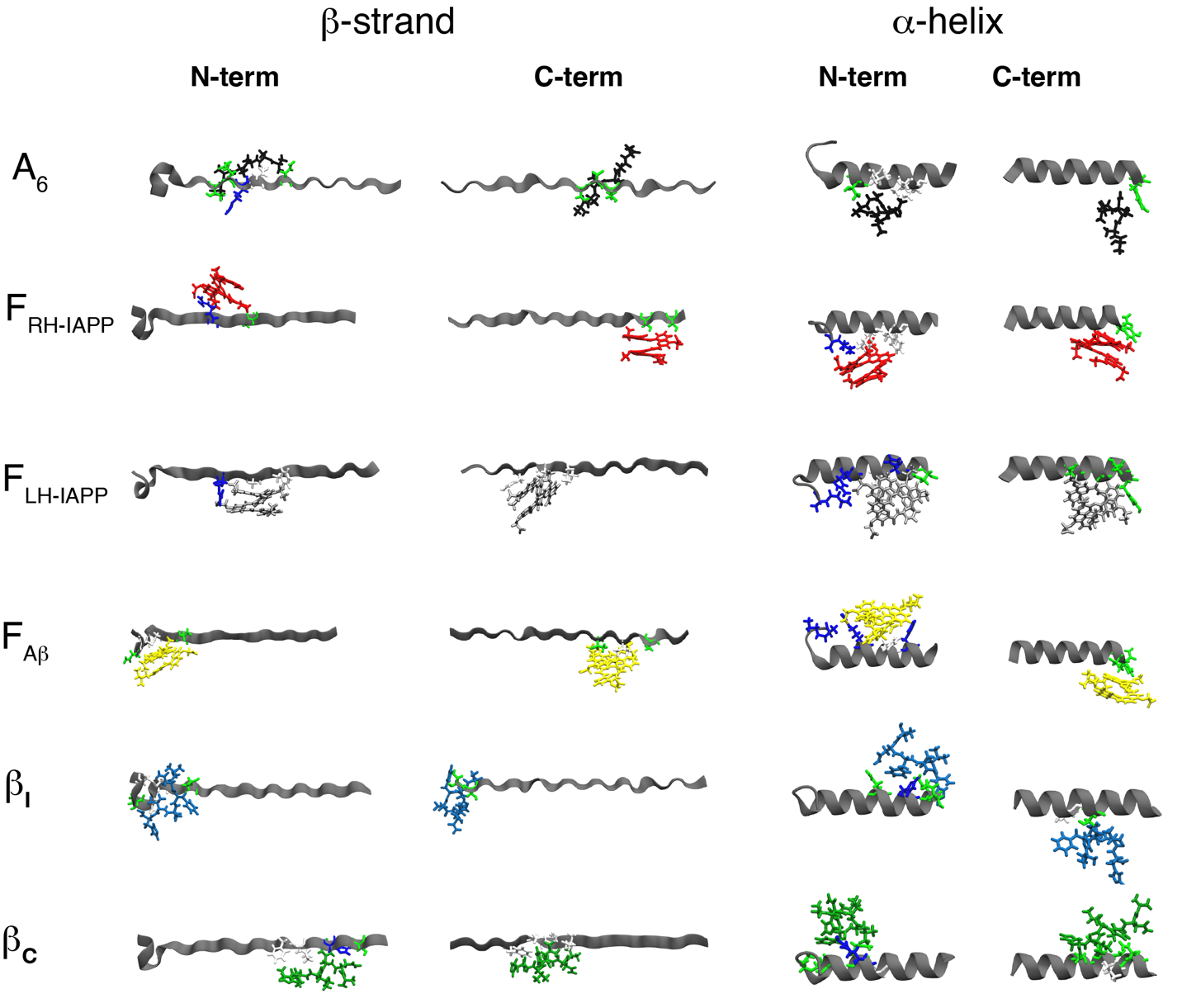

**Figure S3.** Representative complexes after clustering binding simulations for each binder to *human* IAPP in either β-strand or ⍺-helix (grey ribbons). Each inhibitor is distinguished by color: PolyA (A_6_, black); RH-IAPP Foldamer (F_RH-IAPP_, red); LH-IAPP Foldamer (F_LH-IAPP_, grey); Aβ-Foldamer (F_Aβ_, yellow); β-inhibitor (β_I_, blue); and control peptide (β_C_, green). Residues within 4 Å of inhibitor shown (colored by residue type: hydrophobic, white; polar, green; positive charged, blue).

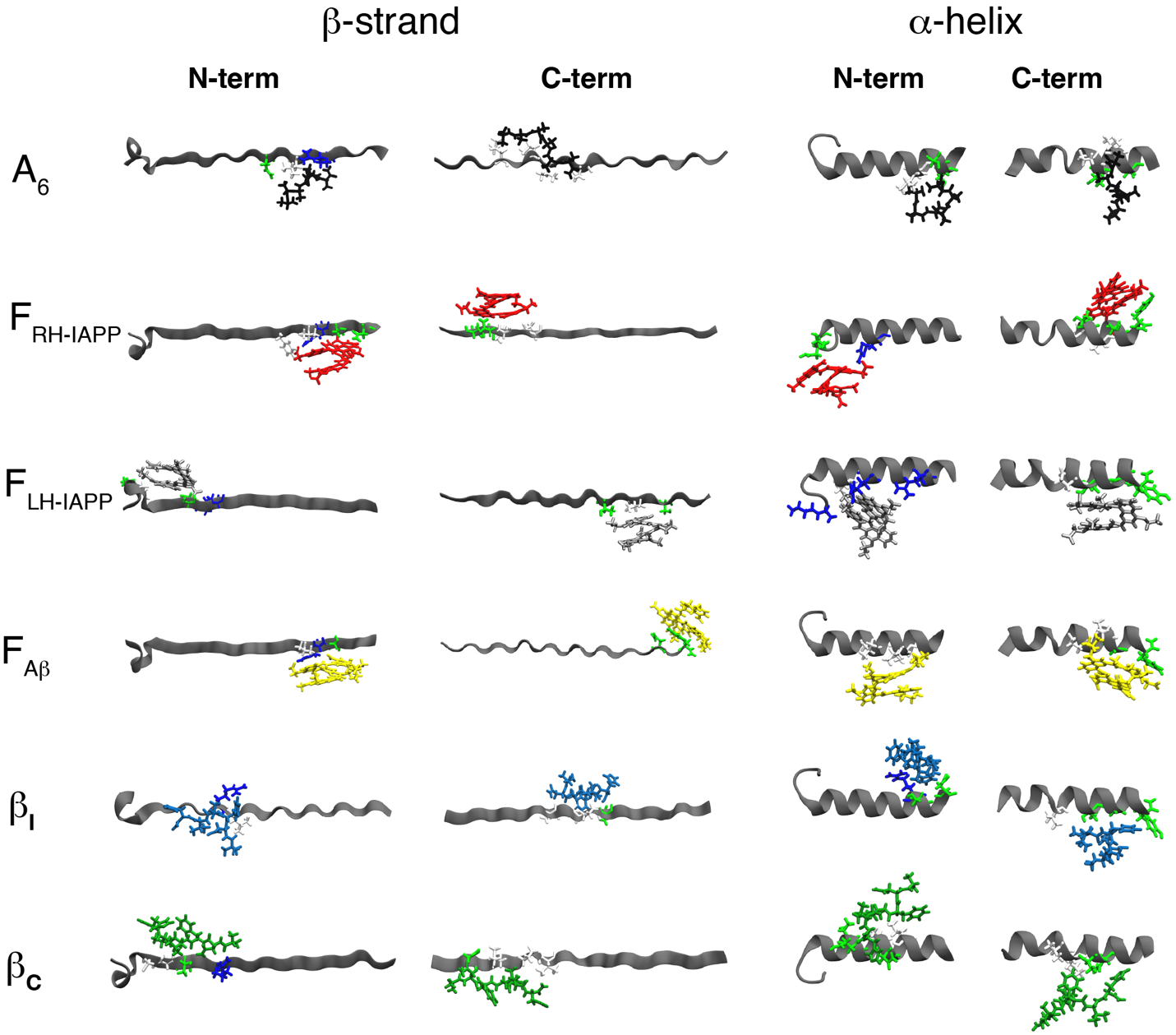

**Figure S4.** Representative complexes after clustering binding simulations for each binder to *rat* IAPP in either β-strand or ⍺-helix (grey ribbons). Each inhibitor is distinguished by color: PolyA (A_6_, black); RH-IAPP Foldamer (F_RH-IAPP_, red); LH-IAPP Foldamer (F_LH-IAPP_, grey); Aβ-Foldamer (F_Aβ_, yellow); β-inhibitor (β_I_, blue); and control peptide (β_C_, green). Residues within 4 Å of inhibitor shown (colored by residue type: hydrophobic, white; polar, green; positive charged, blue).

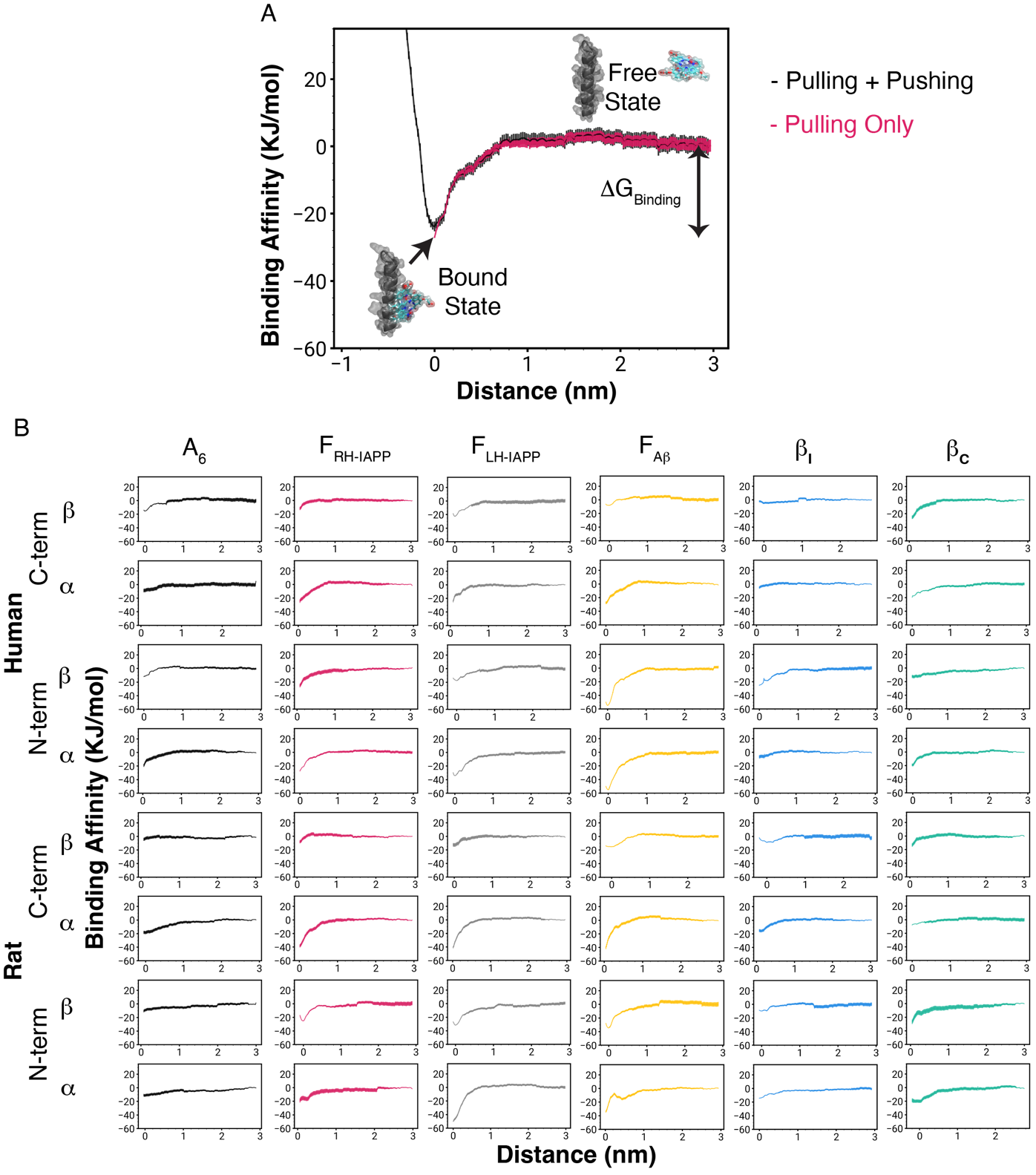
**Figure S5. (A)** Representative potential of mean force curve from umbrella simulations. Pushing or pulling the inhibitor from equilibrium results in an increase in free energy, confirming the lowest energy complex was obtained. The binding affinity (ΔG) is calculated by taking the difference between the “bound” and “free” states. **(B)** Potential of Mean Force plots for all the final complexes simulated in this study: PolyA (A_6_, black); RH-IAPP Foldamer (F_RH-IAPP_, red); LH-IAPP Foldamer (F_LH-IAPP_, grey); Aβ-Foldamer (F_Aβ_, yellow); β-inhibitor (β_I_, blue); and control peptide (β_C_, green). Energy in units of KJ/mol and distance is given in units of nm.

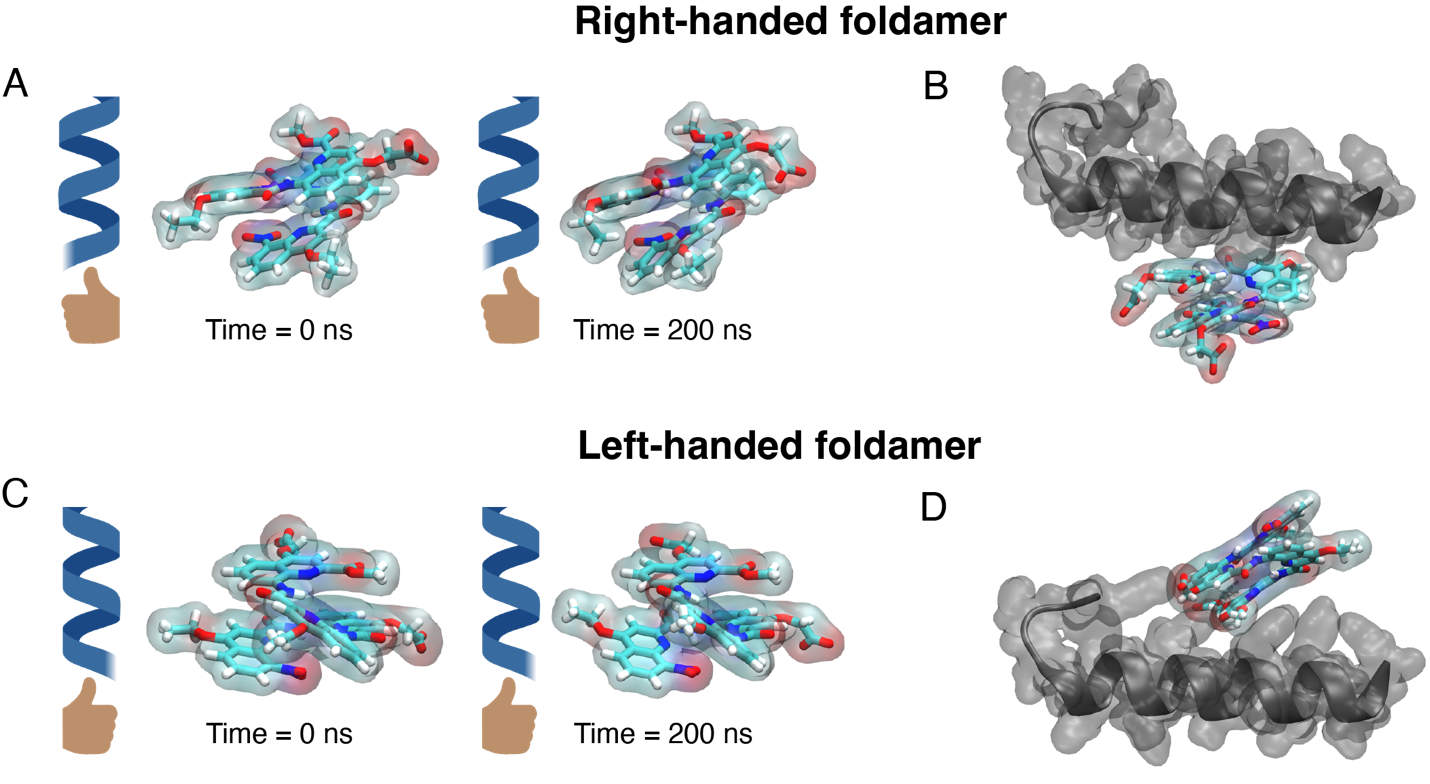

**Figure S6.** Close-ups of **(A)** right-handed and **(C)** left-handed IAPP foldamer at the start (0ns) or end (200ns) of the binding simulation. Foldamers do not interconvert during 200 ns simulations. **(B)** Right-handed or **(D)** left-handed foldamer in complex with helical IAPP (grey) at 200ns (colored by atom type: H, white; C, Teal; N, Blue; O, Red).

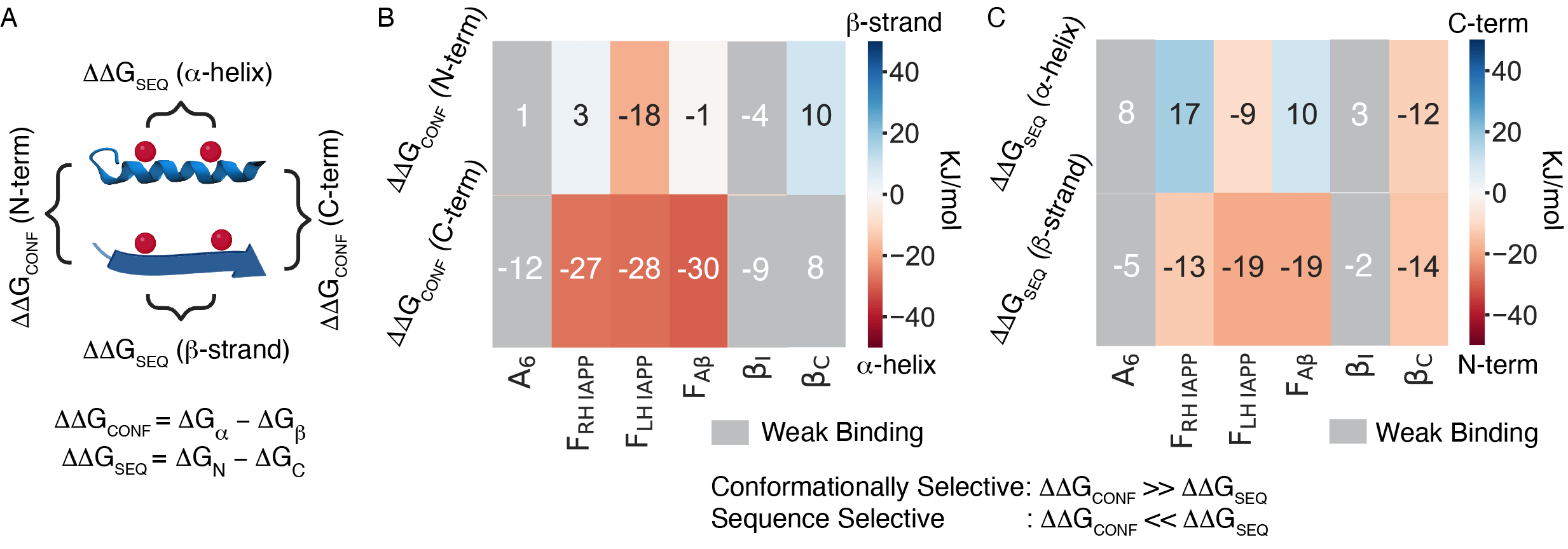

**Figure S7.** **(A)** Conformation and sequence selection for each binder calculated from differences in ΔG of binding (ΔΔG) to *rat* IAPP. **(B)** Conformational selection of IAPP binders, where red are binders with ⍺-helical selection and blue are binders with β-sheet selection. **(C)** Sequence-selection of binders, where red are binders with N-terminal selection and blue are binders with C-terminal selection. Energy units in KJ/mol.

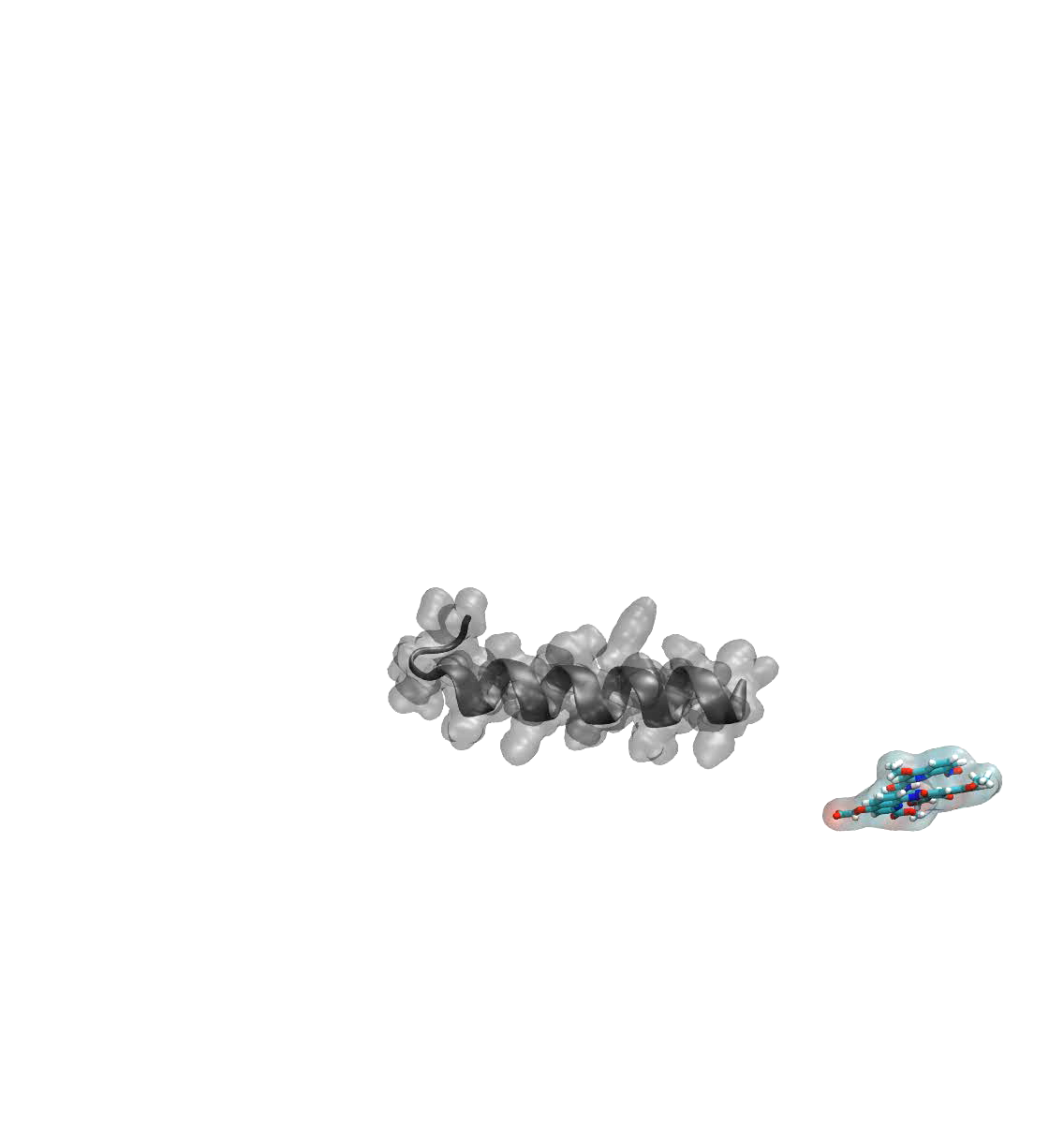

**Movie 1.** Representative converged simulation. Representative simulation of the IAPP-Foldamer binding IAPP within the 200 ns of simulation time.

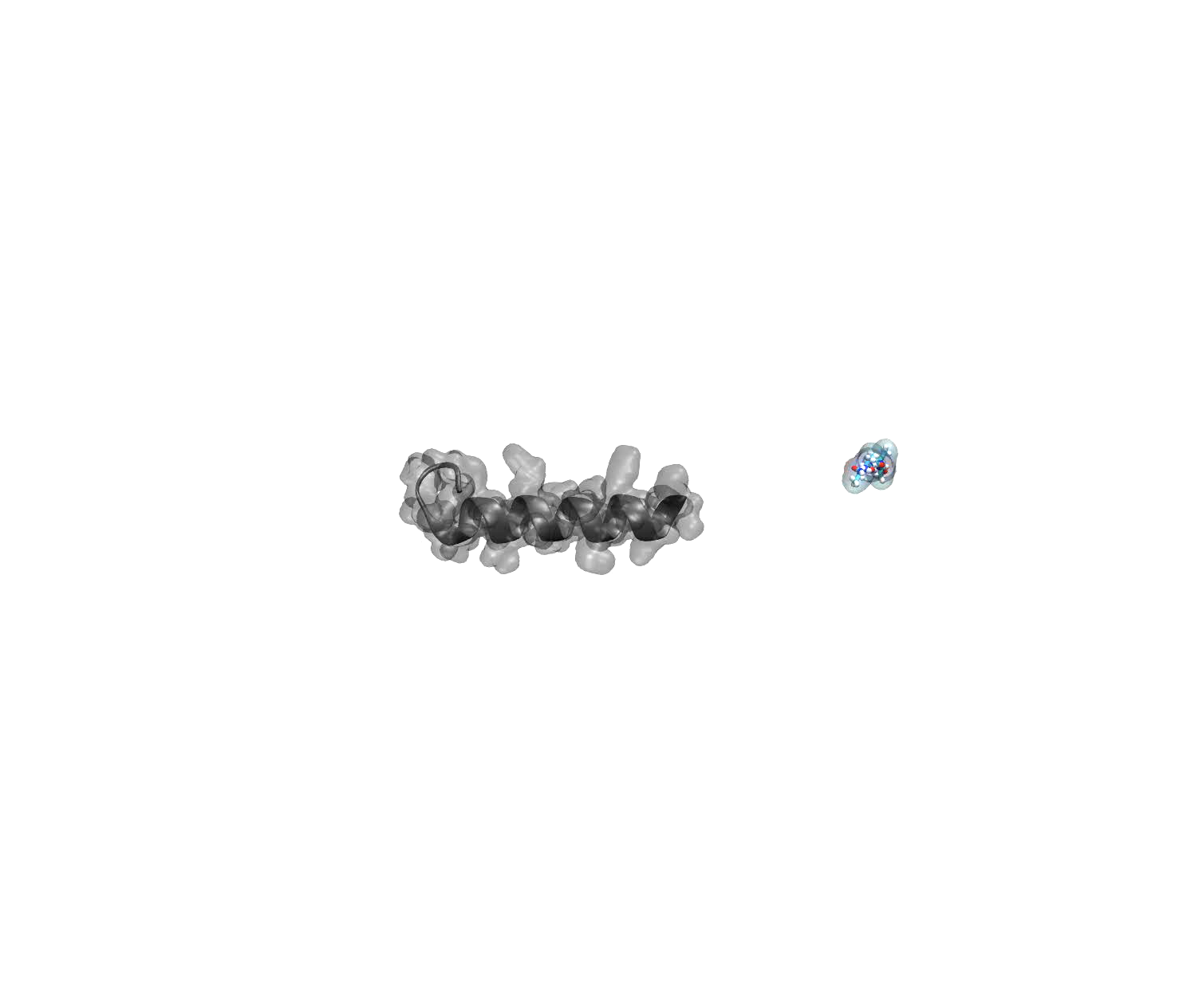

**Movie 2.** Representative non-converged simulation. Representative PolyA simulation does not stably bind IAPP during the 200 ns simulation.

**TABLES**

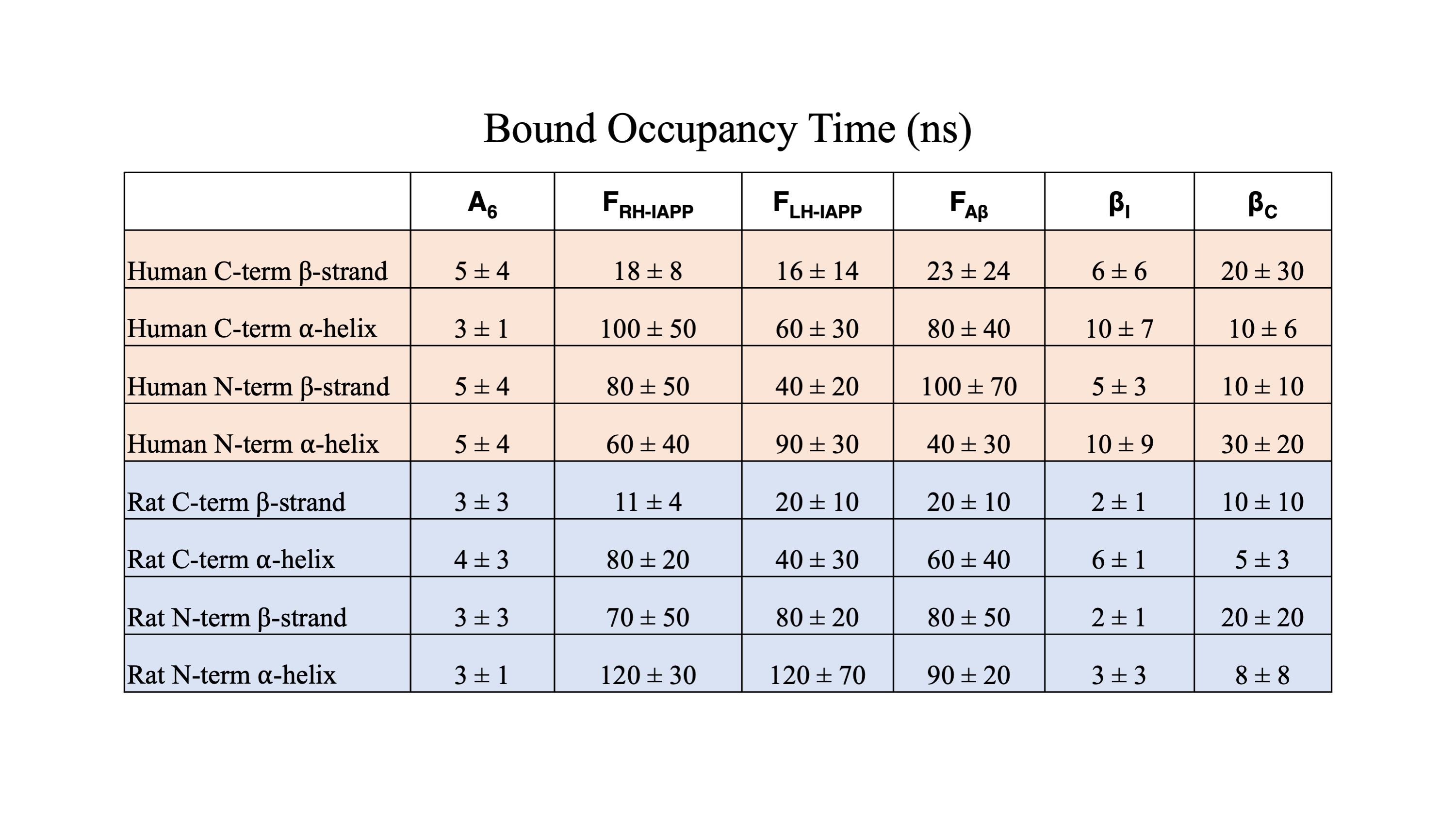

**Table 1.** Bound occupancy time of IAPP binders: PolyA control (A_6_), RH-IAPP Foldamer (F_RH-IAPP_), LH-IAPP Foldamer (F_RH-IAPP_), Aβ foldamer (F_Aβ_), β-inhibitor (β_I_), and control peptide (β_C_) for (peach) human or (blue) rat IAPP with fixed secondary-structure restrained in either β-strand or ⍺-helical conformation over 200ns simulation (in units of ns, N=4).

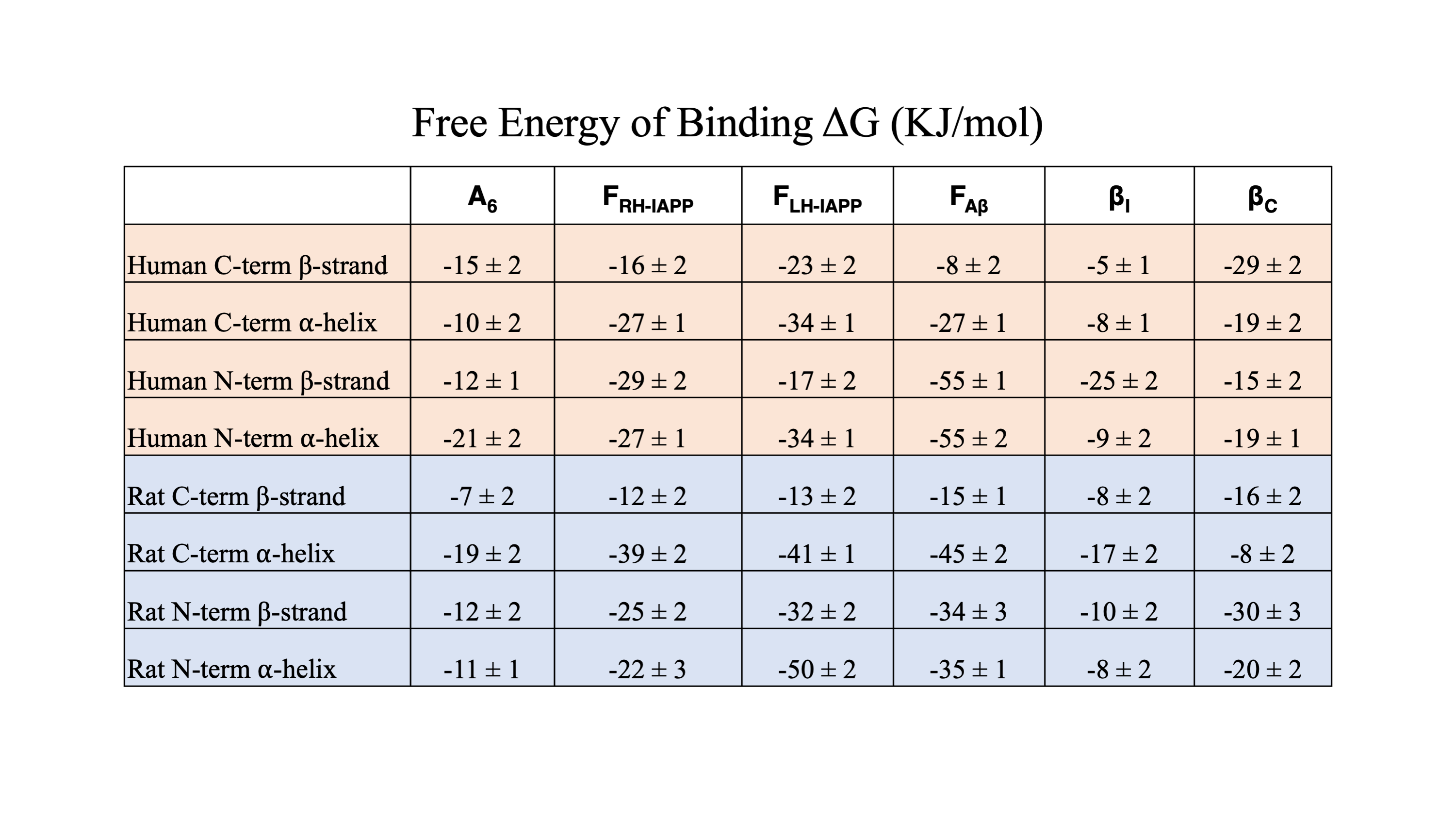

**Table 2.** Binding Affinities of IAPP binders: PolyA control (A_6_), RH-IAPP Foldamer (F_RH-IAPP_), LH-IAPP Foldamer (F_RH-IAPP_), Aβ foldamer (F_Aβ_), β-inhibitor (β_I_), and control peptide (β_C_) with (peach) human or (blue) rat IAPP with fixed secondary-structure restrained in either β-strand or ⍺-helical conformation (in units of KJ/mol).

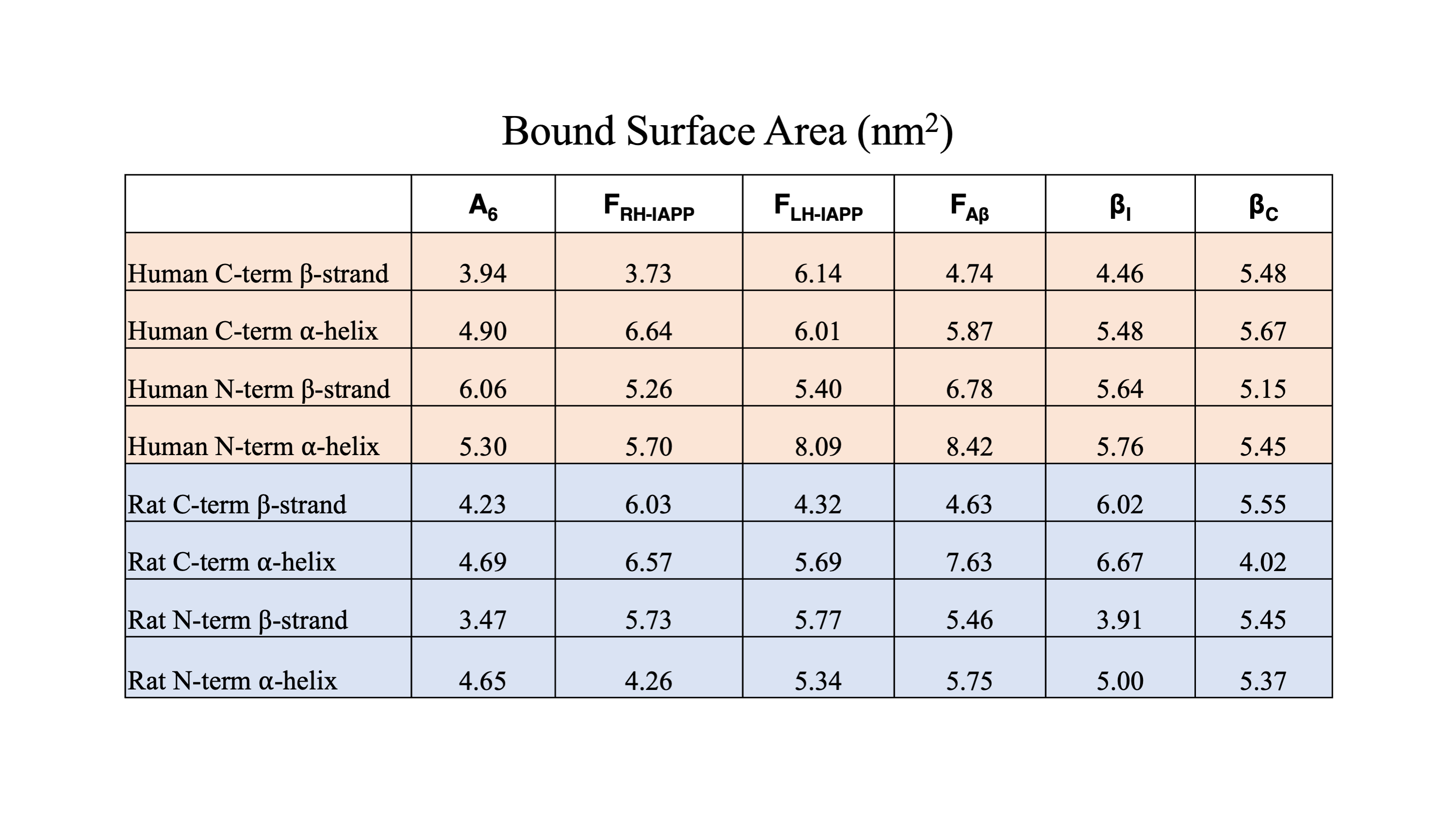

**Table 3.** Bound surface area (BSA) of IAPP binders: PolyA control (A_6_), RH-IAPP Foldamer (F_RH-IAPP_), LH-IAPP Foldamer (F_RH-IAPP_), Aβ foldamer (F_Aβ_), β-inhibitor (β_I_), and control peptide (β_C_) with (peach) human or (blue) rat IAPP in a fixed conformation (in units of nm^2^). (BSA = SASA_IAPP_ + SASA_Inhibitor_ – SASA_Complex_). Larger BSA values represent a larger binding interface between the inhibitor and IAPP.

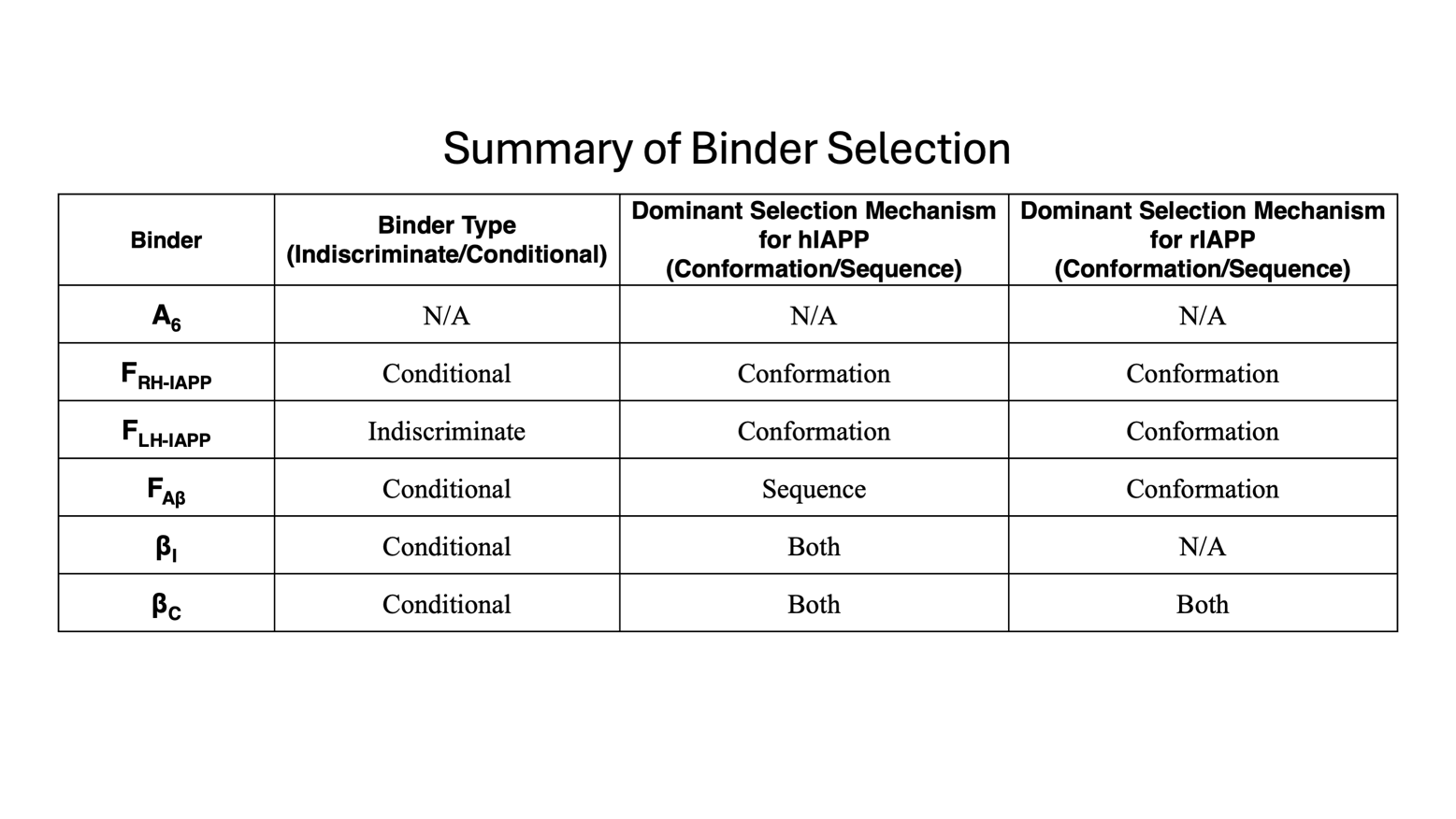
**Table 4.** Summary of selectivity and mechanisms of binders in this study to hIAPP and rIAPP. Indiscriminate binders retain conformational selection despite changes in sequence, while conditional binders only have conformational selection for a particular sequence. Conformation-specific binders bind a particular conformation, irrespective of sequence. Sequence-specific binders bind a particular sequence, irrespective of conformation. Sometimes binders have both.
